# Rapid convergence toward an agriculturalist regulatory landscape following lifestyle transition in Bornean hunter-gatherers

**DOI:** 10.64898/2026.08.09.743820

**Authors:** Pradiptajati Kusuma, Guy S. Jacobs, Hirzi Luqman, Firda A. Ma’ruf, Prisca C. Limardi, Herawati Sudoyo, J. Stephen Lansing, Safarina G. Malik, Irene Gallego Romero

**Affiliations:** Genome Diversity and Disease Division, Mochtar Riady Institute for Nanotechnology, Universitas Pelita Harapan, Indonesia; Human Genomics and Evolution Laboratory, St. Vincent’s Institute of Medical Research, Australia; Department of Archaeology, University of Cambridge, UK; Santa Fe Institute, USA; Centre for Genomics, Evolution and Medicine, Institute of Genomics, University of Tartu, Estonia; School of Medicine, Faculty of Medicine, Dentistry and Health Sciences, University of Melbourne, Australia; Barts Cancer Institute, Queen Mary University of London, UK

**Keywords:** Borneo, Punan, hunter-gatherer, lifestyle transition, genomics, transcriptomics, methylation

## Abstract

The transition from hunter-gatherer to agricultural lifestyles is one of the landmarks in the history of human societies, but its phenotypic and immediate molecular consequences remain poorly understood. The Punan of Borneo clarify this question: two closely related groups — the Punan Batu, still practising mobile foraging, and the Punan Tubu, resettled and farming within the last three generations — share deep common ancestry and no detectable post-split gene flow from agricultural neighbours, allowing the molecular consequences of lifestyle change to be separated from underlying genetic architecture. We applied a multi-omics approach across these two groups and a neighbouring agriculturalist community, the Lundayeh, profiling whole-genome sequences, blood transcriptomes, and DNA methylation arrays. Despite their shared ancestry, the resettled Punan exhibit transcriptomic and methylation profiles more similar to the agriculturalists than to their still-foraging Punan relatives, with 87% fewer differentially expressed genes and substantially fewer differentially methylated sites in the Punan Tubu versus Lundayeh comparison, and show a 77% shift towards an agricultural state. Genes associated with fat metabolism and immune function are the main components of this molecular convergence. These findings demonstrate that lifestyle transition can reshape the human transcriptome and DNA methylation on timescales far shorter than genomic evolution, with implications for understanding the biological consequences of rapid subsistence change in Indigenous communities worldwide.

## Introduction

The biological implications of lifestyle transition are a subject of human evolutionary interest: a transition experienced by the ancestors of the vast majority of the global population, in various places and at varying times (Luca et al., 2010; Larsen, 2023). Anthropological and archaeological research has documented hunter-gatherer lifeways (Binford, 1990; Winterhalder, 2001; Bowen and Gleeson, 2019), while work in behavioural ecology has studied hunter-gatherer subsistence economies (Marlowe, 2005; Kelly, 2013), life histories (Gurven and Kaplan, 2007), and cooperative practices (Apicella et al., 2012) among diverse subjects. Biological and genetic research has explored differences in metabolism and immunity (Pontzer et al., 2012; Urlacher et al., 2018), the microbiome (Schnorr et al., 2014; Smits et al., 2017), and signatures of selection (Fumagalli et al., 2015; Lopez et al., 2019). Yet, integrative perspectives on the nature of transition are lacking — and with it the causal chain from the experience of transition to longer-term evolutionary outcomes. In this study, we ask how gene regulation shifts with when hunter-gatherers adopt agricultural life and how this can inform understanding of natural selection during ancestral transitions.

In Borneo, the Punan Batu hunter-gatherer community, one of a vanishingly small number of groups worldwide still occupying karstic rock shelters, lives in a network of shifting camps in the forest. Punan Batu people continue to hunt daily and typically move location every 8-9 days (Lansing et al., 2022). Genetic evidence suggests the Punan diverged from proto-Austronesian populations before the Austronesian expansion into Indonesia and the Pacific (Kusuma et al., 2023), 4,000 to 3,500 years ago (Widianto and Noerwidi, 2023), or later (McColl et al., 2018). The absence of subsequent Austronesian admixture in Punan genomes points to a continuity of their hunter-gatherer ancestry over thousands of years (Kusuma et al., 2023).

Most Punan groups have resettled and shifted towards a sedentary, agricultural way of life (Sellato, 1994). The resettlement of the Punan Tubu community occurred in the early 1970s (Dounias et al., 2004), when they were relocated by the government from the upstream Tubu River to a resettlement village near the city of Malinau, North Kalimantan (Levang et al., 2005). In under five generations, they abandoned their foraging lifestyle, becoming rice cultivators whose economy now depends on market activities; timber and coal concession fees; and urban amenities, including electricity, schools, and healthcare (Dounias et al., 2007). What makes this transition interesting as a study system is that their closely related population — the Punan Batu — is still largely practising the hunter-gatherer lifestyle, therefore the lifestyle transition can be studied by comparing them against the genetic and regulatory baseline of their contemporary hunter-gatherer relatives.

Genomic evolution operates over thousands of generations (Karlsson et al., 2014; Akbari et al., 2026), far too slow to account for the biological consequences of a transition that occurred within living memory. Yet, populations that shift from foraging to agriculture face fundamentally different dietary intakes, activity levels, pathogen exposures, and metabolic demands (Campbell and Ranciaro, 2021; Malmström et al., 2010; Shah et al., 2022; Harrison et al., 2019). Whole genome sequencing studies have shown that hunter-gatherer populations exhibit genetic adaptations related to foraging diets, environmental conditions, and immunity (Hsieh et al., 2016; Lopez et al., 2019; Herzog et al., 2025), including variations in genes associated with metabolism and nutrient absorption (Zhang et al., 2021), raising the question of how populations adapt. Gene expression is highly sensitive to environmental change (Hamann et al., 2020; Ballinger et al., 2023), and DNA methylation can rapidly shift in response to dietary or environmental perturbation (Fagny et al., 2015; Hernando-Herraez et al., 2015; Zhang and Kutateladze, 2018). This suggests a multi-layered biological response: regulatory rather than genomic adaptation over shorter timescales, followed by longer-term genetic adaptation, an evolutionary interplay between regulatory plasticity, fitness and genetic variation. The juxtaposition of genomic, transcriptomic, and epigenetic data, and the contrast in their evolutionary rates of response (Blake et al., 2020; Bogan and Yi, 2024; Ghalambor et al., 2015), therefore provides the appropriate approach for asking how fast, how far, and through which molecular mechanisms a population can shift its biology when its lifestyle changes.

Here, we describe genomic, transcriptomic, and regulatory dimensions from whole blood sample of lifestyle transition in three Bornean populations: the Punan Batu, hunter-gatherers to this day; the Punan Tubu, resettled former hunter-gatherers; and the Lundayeh, an Indigenous agriculturalist group living in proximity to the Punan Tubu resettlement village (**Fig. 1A**). Our findings indicate that phenotypic shifts in metabolic and fat-related traits during agricultural transition are primarily driven by changes in gene expression and epigenetic regulation. By applying a multi-omics approach to studying these groups as a model, we can better understand the evolutionary impact of lifestyle changes on diverse Indigenous populations around the world.

**Figure 1.**
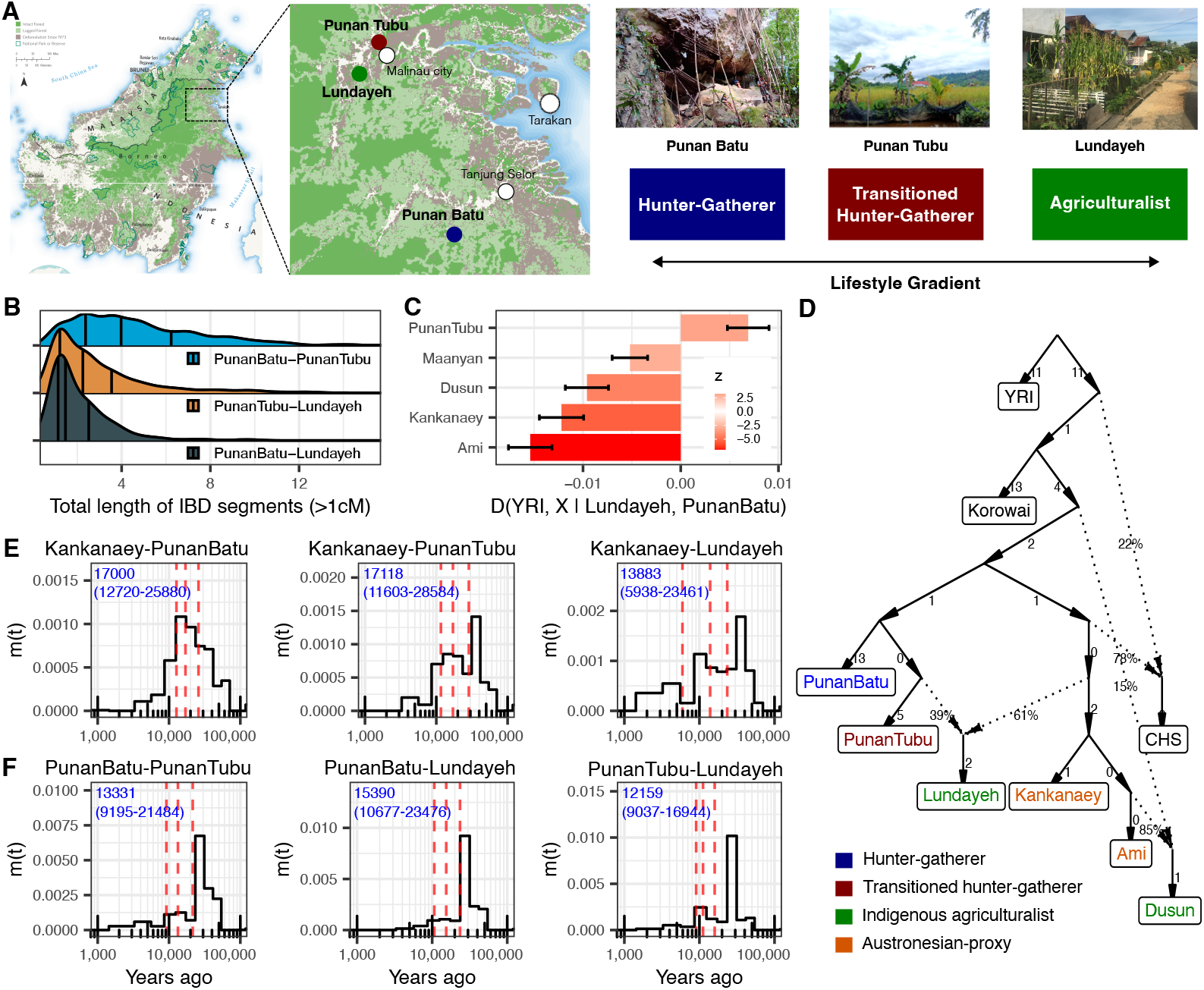
Population history and shared ancestry of Bornean hunter-gatherer and agriculturalist groups. (**A**) Geographic locations of the three study populations in Borneo (cartography by Chris Bruce/TNC (The Nature Conservancy, 2023)) and a schematic of the lifestyle transition gradient, with photographs representative of each site shown. (**B**) Interpopulation IBD comparisons across all pairwise group combinations. (**C**) D-statistics `[D(YRI, X | Lundayeh, Punan Batu)]` quantifying allele-sharing patterns across comparative populations; positive values indicate greater affinity to Punan Batu, negative values indicate greater affinity to Lundayeh. (**D**) Admixture graph (`qpGraph;` worst *Z*-score = 3.008) depicting the inferred population topology and admixture history. (**E**) Cross-coalescence rates from `MSMC-IM` between the Kankanaey (Austronesian proxy) and each of the three study populations, estimating the timing of divergence between Austronesian-related ancestry and the Bornean groups. (**F**) Cross-coalescence rates from `MSMC-IM` among the three study populations — Punan Batu vs. Punan Tubu, Punan Batu vs. Lundayeh, and Punan Tubu vs. Lundayeh — estimating within-Borneo divergence times.

## Results

### The Punan share a common genetic ancestry distinct from the Lundayeh

To establish the demographic baseline for interpreting molecular differences between groups, we performed genetic analyses from whole genome sequence data from the three populations, together with comparative populations from public and published datasets (**Table S1**). Principal component analysis confirmed that Punan individuals form a cluster associated with, but distinct from, other Bornean groups in our dataset (**Fig. S1A**). Punan Batu carry long Runs of Homozygosity (RoH), at the same level as another hunter-gatherer group in the region, the Orang Rimba from Sumatra, and longer than the Jarawa/Onge in the Andaman (**Fig. S1B**), consistent with the small effective population size (**Fig. S1C**).

The interpopulation Identity by Descent (IBD) analysis revealed considerably longer cumulative shared IBD fragments between Punan Batu and Punan Tubu (mean = 4.628 cM, median = 3.975 cM) compared to all other pairwise comparisons (Kruskal-Wallis *p <* 2 × 10 ^− 16^) (**Fig. 1B**). Similarly, D-statistics also show significant allele frequency correlations between the Punan Batu and Punan Tubu compared to between either group and the Lundayeh, other agriculturalists, or Austronesian proxies such as the Kankanaey and Ami (Larena et al., 2021) (**Fig. 1C**), supporting genetic relatedness between the Punan groups that is i) driven by demographic history rather than geography, given similar locations of the Punan Tubu and Lundayeh and ii) has not been disrupted by any recent admixture into our Punan Tubu sample during transition.

Indeed, these ancestral connections are deep. Admixture graph modelling (qpGraph) placed the two Punan groups in a separate clade distinct from Austronesian-related proxies (**Fig. 1D**), unlike the Lundayeh, who experienced Austronesian-related admixture. Cross-coalescence analysis using MSMC-IM (Wang et al., 2020) estimated the split between Austronesian-related groups and the Punan groups at approximately 17,000 years ago (**Fig 1E**), well before the Austronesian dispersal (Denham, 2014; Skoglund et al., 2016), with no evidence of subsequent gene flow from Austronesian groups into either Punan group (**Fig. 1D,E**). Cross-coalescence between the Punan groups and Northeast Bornean agriculturalists (Lundayeh) indicated similarly ancient splits — approximately 15,000 (10,000-23,000) years ago for Punan Batu and 12,000 (9,000-17,000) years ago for Punan Tubu — with little to no gene flow thereafter (**Fig. 1F**). An Austronesian-related pulse occurred into the Lundayeh around 5,000 years ago (**Fig. 1E**) but is not observed among the Punan. These analyses establish the foundation of our evolutionary analyses, demonstrating deep splits between communities that will have allowed for genetic divergence and independent selection over many millennia, including divergent lifeways over the last 5,000 years.

### Resettled Punan occupy an intermediate phenotypic position between hunter-gatherers and agriculturalists

With a genetic demographic scaffold in place, we now turn to the phenotypic manifestation of lifestyle transition. We compared a range of anthropometric, body composition, and metabolic traits across the three groups. Both Punan Batu and Punan Tubu were of shorter stature compared to the Lundayeh, with the average male height across the Punan communities being 156 cm (**Fig. 2A**), 6 cm shorter than the Lundayeh average and 3 cm taller than the Batwa hunter-gatherer community in Africa (Perry et al., 2014). Punan Batu individuals show the most distinct body composition, with lower body weight, waist and hip circumferences, and consequently lower BMI compared to the other groups, as well as lower fat mass but higher muscle mass on bioimpedance measurements (**Fig. 2B**). Blood biochemistry similarly distinguishes Punan Batu, with lower glucose, cholesterol, and triglyceride levels (**Fig. 2C**). These differences are consistent with their dietary patterns. Based on food-frequency questionnaires and direct observations, Punan Batu consume significantly more tubers and red meat and markedly less rice than the other groups (**Fig. 2D**), the latter reflecting a community taboo against rice cultivation, with rice instead obtained through trade.

**Figure 2.**
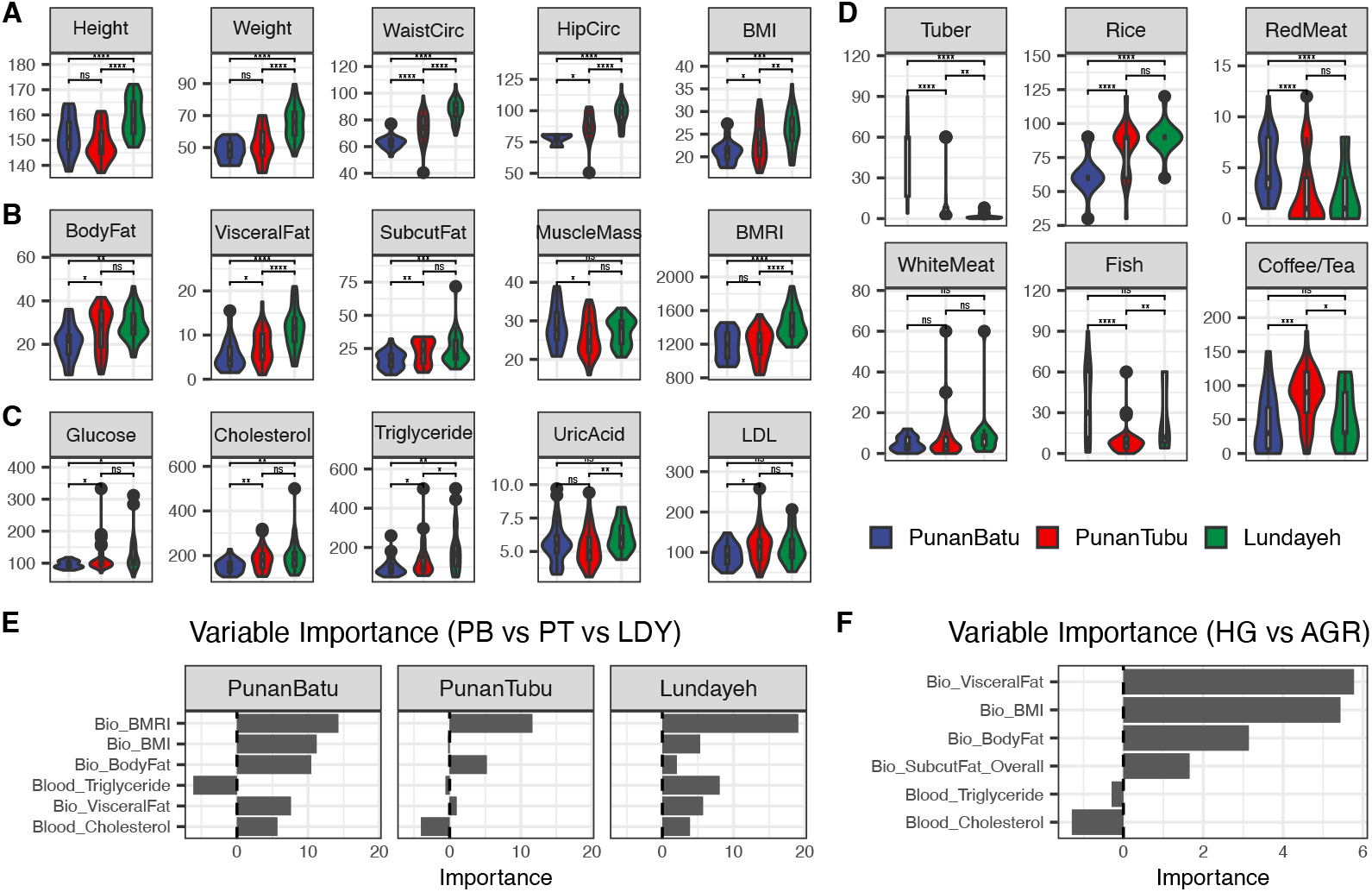
Phenotypic intermediacy of the resettled Punan across traits and diets. (**A**) Anthropometric measurements across Punan Batu (PB), Punan Tubu (PT), and Lundayeh (LDY). (**B**) Body composition measured by bioimpedance analysis. (**C**) Blood biochemistry from point-of-care testing. (**D**) Dietary frequency across food categories. (**E**) Permutation-based variable importance from the ternary Random Forest classifier (PB vs PT vs LDY). (**F**) Permutation-based variable importance from the binary Random Forest classifier (hunter-gatherer versus transitioned).

Notably, Punan Tubu individuals do not resemble one group or the other; they often occupy an intermediate phenotypic position. To quantify this, we tested 3 models on six phenotypic measures that significantly differ across groups; Kruskal-Wallis FDR-adjusted *p ≤* 0.05, highest-ranked effect sizes; **Table S3**) to determine which phenotypes are most predictive of population group and hence lifestyle. Among those models, the Random Forest gained better accuracy (**Table S4**). Basal metabolic rate index was inferred as the strongest predictor of lifestyle, followed by body fat and BMI (**Fig. 2E**). Model performance was highest in classification of Punan Batu (AUC = 0.81), followed by Lundayeh (AUC = 0.80) and Punan Tubu (AUC = 0.67) (**Fig. S2A**,**C**), reflecting the intermediate and overlapping phenotypes of the Resettled Punan. To more directly characterise the direction of this transition, we recoded the outcome into a binary variable: hunter-gatherer (Punan Batu) versus transitioned (Punan Tubu and Lundayeh combined). Model performance improved from 56% accuracy *(κ* = 0.34) to 77% (*κ* = 0.40) (**Table S4**) with ROC curve AUC at 0.79 (**Fig. S2B**,**D**) under this binary framework, with RF sensitivity of 53% for the hunter-gatherer class and a specificity of 88% for the transitioned class. Variable importance again highlighted BMI and visceral fat as primary drivers, with total and subcutaneous body fat contributing moderately (**Fig. 2F**). The phenotypic picture that emerges is therefore a directional transition, such that Punan Tubu have shifted away from the hunter-gatherer phenotype, with body composition, particularly fat distribution, as the main axis of change.

### Resettled Punan’s transcriptome and methylome have converged toward the agriculturalist

The genetic relationship between populations clusters the Punan Batu and Punan Tubu communities, while phenotypic variation influenced by development and dietary practices places the Punan Tubu as intermediate with similarities to long-term agriculturalists. As gene expression is highly plastic, we hypothesise that regulatory variation might also place the Punan Tubu with the agriculturalists, emphasising the scope for rapid biological response to shifting environments and lifestyle. We performed principal component analysis (PCA) and differential analysis on TMM-normalised RNA sequencing data (13,710 genes, *Data S1)* and pre-processed DNA methylation array data (880,992 CpGs, *Data S2)* across all three groups (see Methods) to observe these. In contrast to the separation of all three groups in genomic PCA (**Fig. 3A**), Punan Tubu visibly clusters with Lundayeh in the RNA seq and methylation PCAs (**Fig. 3B-C**).

**Figure 3.**
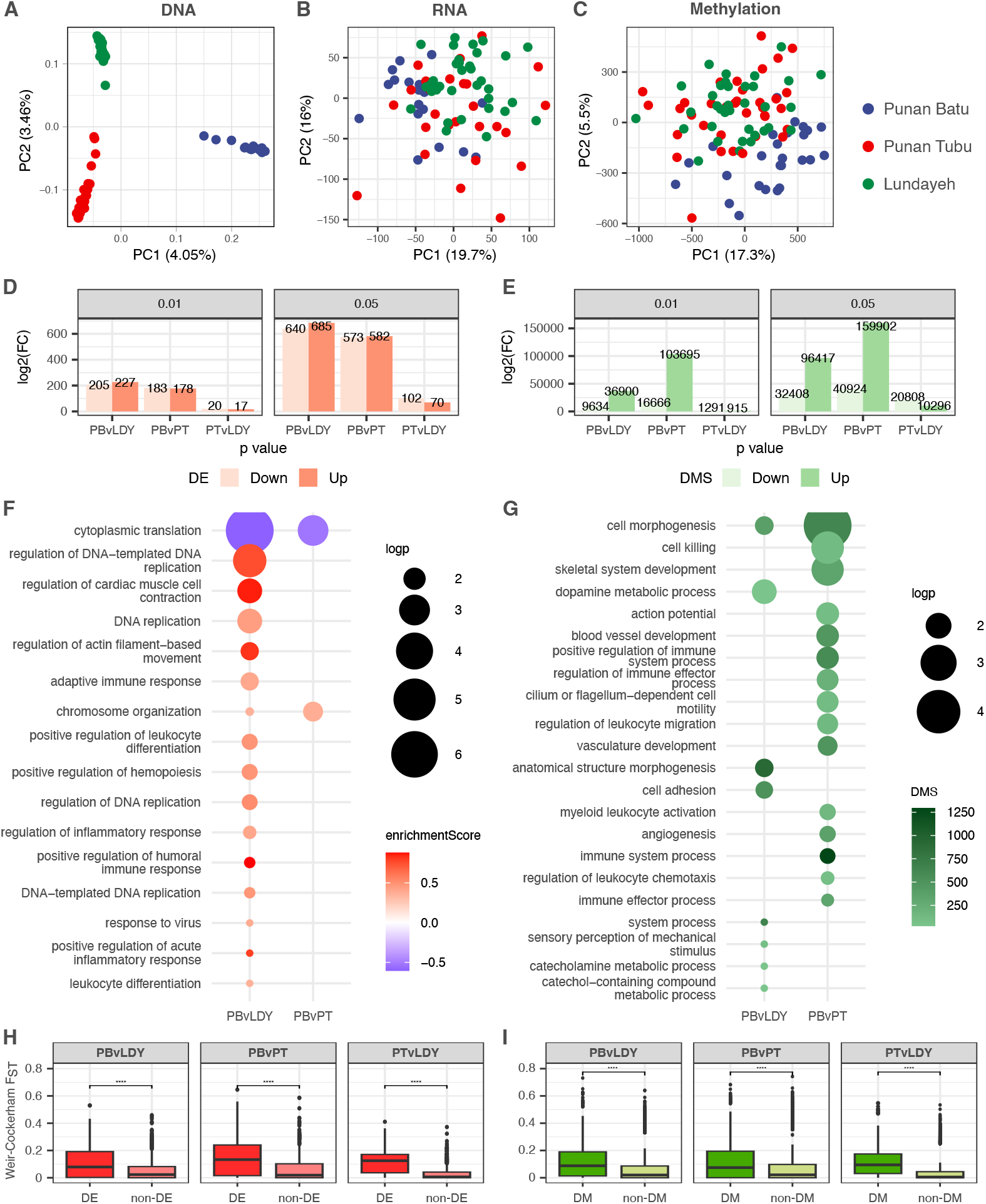
Convergence of transcriptomic and DNA methylation profiles in the resettled Punan toward the Lundayeh agriculturalist. (**A-C**) Principal component analysis of genomic DNA (**A**), RNA-seq (**B**), and DNA methylation (**C**) data across Punan Batu hunter-gatherer (PB), Punan Tubu resettled hunter-gatherer (PT), and Lundayeh agriculturalist (LDY) populations. (**D-E**) Numberof differentially expressed genes (DEGs; **D**) and differentially methylated sites (DMS; **E**) across the three pairwise comparisons at varying FDR thresholds. (**F-G**) Functional enrichment of significant DEGs by GSEA and GO Biological Process analysis (**F**) and of significant DMS by `GOmeth` over-representation analysis (**G**). (**H-I**) Cis-driven DEGs (**H**) and DMSs (**I**) are associated with eQTLs and methylQTLs carrying larger fixation indices (F_ST_) between populations.

Pairwise differential analyses make this convergence quantitatively clearer. We identified 1,325 differentially expressed genes (DEGs) between Punan Batu and Lundayeh and 1,155 DEGs between Punan Batu and Punan Tubu, but only 172 DEGs in the comparison between Punan Tubu and Lundayeh (FDR-adjusted *p <* 0.05; **Fig. 3D, Fig. S4**). The same asymmetry holds for DNA methylation, with 165,725 differentially methylated sites (DMS) between Punan Batu and Lundayeh, 200,826 between Punan Batu and Punan Tubu, but only 31,104 between Punan Tubu and Lundayeh (**Fig. 3E**). Despite their deep genetic divergence from the Lundayeh, the resettled Punan Tubu are far more similar to them than to their Punan Batu relatives in gene expression and methylation profiles. Shared DMSs across all three comparisons account for only 1% of all DMS identified (**Fig. S5**), and there are no overlapping DEGs across all three comparisons, confirming that the Punan Tubu and Lundayeh profiles have converged rather than that all three groups are diverging.

Gene set enrichment analysis (GSEA) against Gene Ontology and KEGG pathways (The Gene Ontology Consortium, 2018; Kanehisa and Goto, 2000) identified enriched terms in the Punan Batu versus Lundayeh comparison related to immune response, cytokine production and regulation, and blood circulation (**Fig. 3F, Table S5**). Despite the large number of DEGs in the Punan Batu versus Punan Tubu comparison, only two GO Biological Process terms were enriched, i.e., cytoplasmic translation and chromosome organisation, both of which overlap with the Punan Batu versus Lundayeh comparison, and no significant KEGG pathways were identified. The Punan Tubu versus Lundayeh comparison yielded no significantly enriched terms. Overrepresentation analysis (ORA) produced the same pattern (**Table S6**), showing significant functional enrichment only in the Punan Batu versus Lundayeh contrast, covering immune responses, circulatory processes, and cholesterol storage regulation. Enrichment analysis of DMS similarly identified immunity, neurotransmitter, and cell morphogenesis terms (**Fig. 3G, Table S7**), partially overlapping with the expression enrichment.

To confirm that lifestyle primarily associates with regulatory shifts, we decomposed differential expression and methylation signals by lifestyle (Punan Batu versus Punan Tubu and Lundayeh combined) and by ancestry (both Punan groups combined versus Lundayeh) (**Fig. S6**). The lifestyle comparison largely yielded more DEGs (1,410 genes; 157,879 DMS) than the ancestry comparison (276 genes; 19,327 DMS) at FDR-adjusted *p <* 0.05, confirming that the dominant axis of molecular variation in this dataset aligns with lifestyle rather than genetic ancestry. While the overall log_2_FC directionality trends in the ancestry comparison are broadly concordant, genes and methylation sites where Punan Batu and Punan Tubu diverge in opposing directions relative to Lundayeh (**Fig. S6A**,**G**) identify loci where Punan Tubu has already shifted toward the Lundayeh profile. This pattern is largely absent in the lifestyle comparison, which shows consistent directional concordance across contrasts (**Fig. S6B**,**H**). Functional enrichment of the ancestry comparison identified terms related to lipid metabolism and cholesterol regulation, while the lifestyle comparison captured these same terms alongside additional immune-related processes, such as leukocyte proliferation, lymphocyte proliferation, and T-cell proliferation. Methylation changes in the lifestyle comparison were similarly enriched for immune terms including response to bacterium and type II interferon production, while the ancestry comparison showed no significant methylation enrichment.

To identify the contribution of underlying genetic variation to these molecular differences, we mapped cis expression and methylation quantitative trait loci (eQTL and methylQTL). We detected 914 signfificant cis-eQTLs with 890 unique variants *(Data S3)* and 20,311 significant cis-methylQTLs with 16,917 unique variants *(Data S4)*. Consistent with findings in other population comparisons (Nédélec et al., 2016; O’Neill et al., 2021; Saitou et al., 2024), cis-driven DEGs and DMS have eQTLs and methylQTLs with larger fixation indices (F_ST_) between populations (**Fig. 3H-I**), indicating that a subset of molecular differences is influenced by divergent genetic variants. However, the overall low number of detected QTLs, which is due to limited sample size in our dataset, means that genetic variants explain only a fraction of the observed molecular convergence. These results highlight that the resettled Punan Tubu, to a certain degree, have undergone an apparent transition in their blood transcriptome and DNA methylome converging toward the Lundayeh agriculturalist profile driven by lifestyle.

### Resettled Punan have moved rapidly toward an agriculturalist expression state

To address whether the Punan Tubu convergence towards agriculturalist state is consistent with drift process or directional movement, we modelled gene expression evolution explicitly across the three populations using an Ornstein-Uhlenbeck (OU) framework (Rohlfs et al., 2013) that tests whether expression state has shifted with lifestyle. We fitted five nested models (**Table S8**) per gene across over 13,000 genes using a star trifurcation topology with a shared root age of approximately 9,636 years ago derived from MSMC-IM pairwise split times. This model is a conservative ‘null hypothesis’ in which divergence is largely driven by genetic divergence over the last 10,000 years.

By AIC, BM was the best model for 5,079 genes (37.8%), while OU_*θ*_ for 7,969 genes (59.3%) and OU_shared_ for 382 genes (2.8%) (**Fig. 4A-B**). Likelihood Ratio Tests identified 6,829 genes (50.8%) with significant population-specific *θ* (FDR-adjusted *p <* 0.05). For genes where the LDY-PB expression difference exceeded an arbitrary threshold of 0.1 log_2_-CPM, we defined a transition progress metric *π = (θ*_*PT*_ *— θ* _*PB*_*)/(θ*_*LDY*_ *— θ*_*PB*_), where *π* = 0 and *π* = 1 correspond to the assumed hunter-gatherer and agriculturalist expression stable states, respectively. Among 7,365 genes meeting this threshold, the median transition progress was 0.767 (**Fig. 4D**), indicating that PT has moved approximately 77% of the way from the ancestral hunter-gatherer toward the agriculturalist state. Genes with *π >* 1 (overshoot) and *π <* 0 (opposite direction) were observed in 455 and 70 genes, respectively.

**Figure 4.**
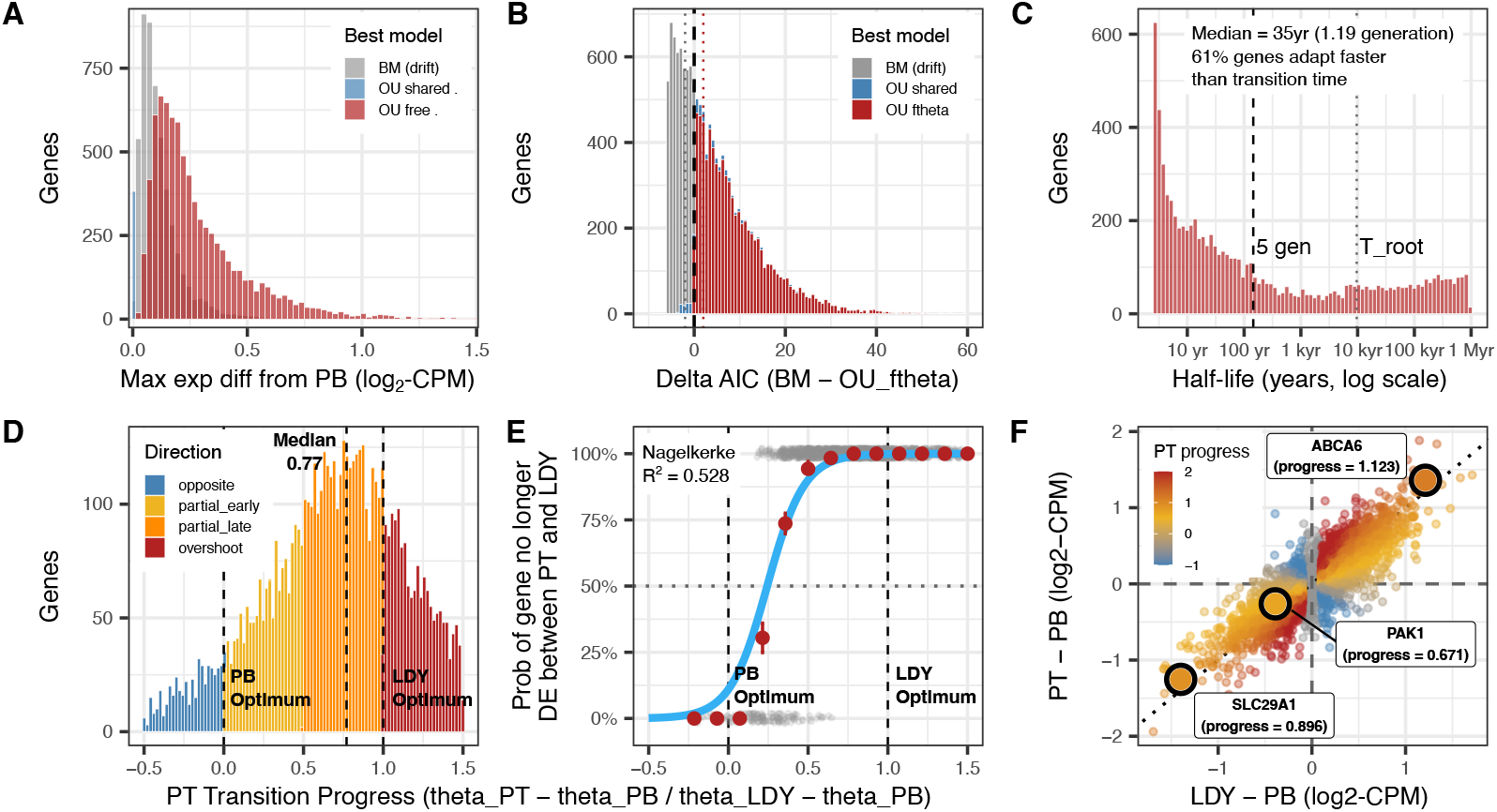
Rapid transcriptomic convergence with the agriculturalist expression state in resettled Punan. (**A**) Maximum absolute expression shift from PB across 13,430 genes, stratified by best-fitting model (AIC). (**B**) Delta AIC distribution (BM **—** OU_θ_) across all genes. (**C**) Half-life distribution ***(t****-*_*1/2*_ = ln_2*/α*_) for OU _θ_ genes. (**D**) PT transition progress *π* = (*θ*_*PT*_ *— θ*_*PB*_*) / (θ*_*LDY*_ *— θ*_*PB*_ ) across 7,365 genes (LDY-PB difference _*>*_ 0.1 log_2_-CPM). (**E**) Probability of DE resolution (gene no longer DE between PT and LDY) as a function of *π*, among genes DE between PB and LDY (FDR _*<*_ 0.05; *n* = 1,897). Logistic regression Nagelkerke pseudo-*R*^*2*^ = 0.528. (**F**) PT vs LDY expression shift from PB for significant OU_θ_ genes (FDR _*<*_ 0.05), coloured by *π*.

The genome-wide median *α* across OU-selected genes was 0.580 per generation, corresponding to a median half-life of 1.19 generations (∼35 years) — the time required to close half the gap between the current expression state and the optimum (**Fig. 4C**). Approximately 61.5% of genes individually have *α* values sufficient to achieve 77% progress within the time frame of lifestyle transition (3-5 gens). Under the median *α*, approximately 94.5% of the shift toward a new optimum would be expected within 5 generations; the observed 77% corresponds to an effective adaptation time of approximately 2.5 generations, consistent with the resettlement timeline. Two lines of evidence indicate that this rapid response reflects transcriptional plasticity. First, the median half-life of 1.19 generations is inconsistent with allele frequency change as the primary driver — even strong positive selection *(s* = 0.1) requires tens to hundreds of generations to sweep standing variation to fixation (Chotai et al., 2025). Second, models allowing population-specific *α* provided no improvement over a shared constraint, whereas under a selection model, *α* reflects the fitness landscape and should differ between subsistence environments. The observed pattern, i.e., population-specific optima (*θ*) with shared constraint strength (*α*), is expected if populations inhabit different transcriptional regimes but respond through the same plastic regulatory mechanisms.

This shift is confirmed by the resolvedness of differential expression. Among the 1,325 genes differentially expressed between PB and LDY, 92.5% are no longer differentially expressed between PT and LDY, despite PT and LDY being genetically distant. This resolvedness rate scales with transition progress. Genes with transition progress *π* between 0.7 and 1.0 show 100% resolvedness, while genes with *π <* 0.2 show only 8.82% (**Fig. 4E**). Logistic regression confirms transition progress as a predictor of DE resolvedness *(β* = 8.59, *z* = 14.49, *p <* 2 × 10^−16^, Nagelkerke pseudo-*R*^*2*^ = 0.582). Three fat-associated genes illustrate the spectrum of transition dynamics (**Fig. 4F**): *ABCA6 (π* = 1.12, overshoot), *SLC29A1 (π* = 0.896, partial transition), and *PAK1 (π* = 0.671, partial transition), spanning the range from full convergence to presumably ongoing shift. These genes are examined in detail in the following section.

These models showed that the resettled Punan Tubu have not drifted randomly away from the hunter-gatherer state, but they have moved directionally toward the agriculturalist expression state driven by transcriptional plasticity. The speed of this shift indicates that the transcriptome and DNA methylation are capable of tracking lifestyle change far faster than genomic evolution alone could produce. Also, for a population undergoing lifestyle transition, the molecular consequences may be largely shifted well before the genomic signature of that transition becomes detectable.

### Immune and metabolic pathways are the primary axes of molecular transition, with selection and expression largely decoupled

Our analyses so far confirm that lifestyle transition is more strongly associated with regulatory shifts than ancestry (Punan vs non-Punan). However, we also identify significant numbers of eQTLs and methylQTLs and find that, at the DNA sequence level, these tend to be diverged between populations. Given thousands of years of genetic divergence between each population pair (**Fig. 1F**), it is possible that regulatory divergence in some genes has been driven by past selection. We therefore calculated a haplotype-based selection score suited to detecting selection over recent timeframes (XP-EHH (Sabeti et al., 2007)) across genomes between population pairwise. To identify gene sets showing common trends among the three populations, we integrated selection, differential expression, and differential methylation signals across all three pairwise contrasts. We applied *k*-means clustering to each gene’s nine vectors of − log_10_(*p*)-values spanning selection (XP-EHH), differential expression, and differential methylation across the three pairwise contrasts: Punan Batu vs. Lundayeh (PBvLDY), Punan Batu vs. Punan Tubu (PBvPT), and Punan Tubu vs. Lundayeh (PTvLDY). The optimal number of clusters (K = 7) was determined by `NbClust` majority rule and validated by silhouette analysis (average width = 0.2) (**Fig. 5A-B**).

**Figure 5.**
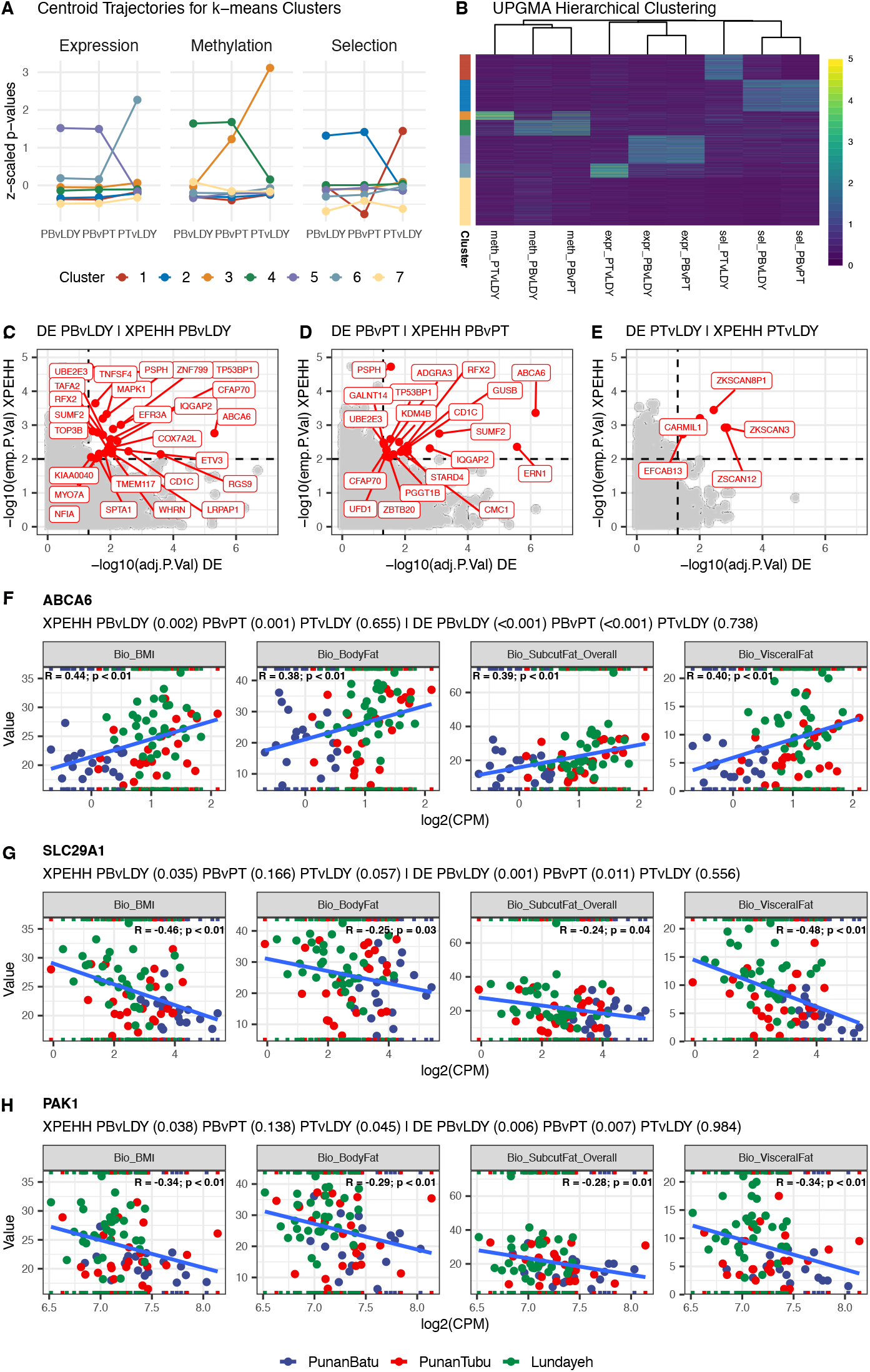
Immune and metabolic pathways drive molecular transition with selection and expression divergence largely decoupled. (**A**) Centroid trajectories for seven *k*-means clusters derived from nine vectors of **-**log_10_ (*p*)-values spanning positive selection (XP-EHH), differential expression, and differential methylation across three pairwise contrasts: PB vs LDY, PB vs PT, and PT vs LDY. (**B**) Heatmap of scaled log_10_ (*p*)-values for all genes grouped by cluster. (**C-E**) Integration of differential expression and XP-EHH selection signals across the PB vs LDY (**C**), PB vs PT (**D**), and PT vs LDY (**E**) contrasts; each point represents a gene, with XP-EHH **-** log_10_ empirical *p*-values plotted against - log_10_ FDR-adjusted *p*-values for differential expression. (**F-H**) Expression levels of three fat metabolism genes — *ABCA6* (**F**), *SLC29A1* (**G**), and *PAK1* (**H**) — across the three population groups (log_2_(CPM)).

The clustering showed overall independent selection, expression, and methylation groups (**Fig. 5B**). There are two clusters, i.e., cluster-5 for expression and cluster-2 for selection, exhibited elevated signals specifically in the PBvLDY and PBvPT contrasts, identifying genes that are distinctively regulated or selected in Punan Batu relative to the other two groups. Two other clusters (cluster-6 and cluster-1) showed elevated signals specific to Punan Tubu contrasts across the same omics layers, capturing genes whose regulatory state is distinctive to the transitional group. Notably, cluster-5, characterised by hunter-gatherer-specific differential expression, showed significant enrichment for body composition phenotypes, such as BMI-associated genes (∼12% of cluster, Fisher’s FDR-adjusted *p* = 0.015) and visceral fat-associated genes (∼17% of cluster, Fisher’s FDR-adjusted *p* = 5.55 × 10 ^9^), representing the only significant phenotype enrichments across all seven clusters (**Fig. S7**). Functional enrichment of cluster-5 further identified KEGG pathways related to lipid metabolism and arrhythmogenic right ventricular cardiomyopathy (FDR-adjusted *p <* 0.01), with suggestive GO Biological Process terms in cholesterol storage and lipoprotein metabolism. These results point to fat metabolism and body composition as the important phenotypes distinguishing hunter-gatherer from post-transition molecular phenotypes, consistent with the phenotypic differences identified before.

Weighted Gene Co-expression Network Analysis (WGCNA) (Langfelder and Horvath, 2008) corroborated this result at the network level. The *red, yellow*, and *black* modules showed positive correlations between module eigengenes and both ancestry and lifestyle and were positively associated with BMI, total body fat, visceral fat, and subcutaneous fat (**Fig. S9**). Conversely, the *magenta, turquoise, light-green*, and *brown* modules showed negative correlations with ancestry and lifestyle and were inversely related to the same fat measures. This co-regulatory structure suggests that Punan Batu harbours a group of co-expressed genes correlated to body composition and that the transition to agriculture is associated with a shift in the expression of these networks. Notably, the *cyan* module showed a strong positive correlation with sex and with subcutaneous and total body fat regardless of lifestyle. This captures a sex-driven adiposity network that works independently of the hunter-gatherer-to-agriculture axis.

The disconnect between selection and expression is a broader pattern, not limited to a cluster of genes. Examining the overlap between significantly differentially expressed genes and the top 1% of XP-EHH scores across all pairwise contrasts (**Fig. 5C-E**), only 0.1% of significant DEGs meet this criterion in the PBvLDY and PBvPT comparisons and just 0.04% in the PTvLDY comparison. Zooming into three fat metabolism genes illustrates how selection, expression, and phenotype intersect across the transition. *ABCA6*, a transporter involved in lipid and cholesterol metabolism (Breuss et al., 2020), shows one of the strongest XP-EHH peaks in Punan Batu relative to both Punan Tubu and Lundayeh, is significantly differentially expressed in those same contrasts (**Fig. 5C-D**), and its transcript levels correlate *(R ≥* 0.38-0.44, FDR-adjusted *p <* 0.01) with BMI, body fat, visceral fat, and subcutaneous fat (**Fig. 5F**). This convergence of selection, expression, and phenotype association suggests that Punan Batu-specific haplotypes at *ABCA6* modulate lipid metabolism that is beneficial under a foraging subsistence strategy. *SLC29A1*, implicated in thermogenic function in human brown adipocytes (Pfeifer et al., 2024), and *PAK1*, involved in adipose tissue metabolism (Rosas and Solaro, 2025), both carry top-5% XP-EHH selection signals in Punan Batu and Punan Tubu relative to Lundayeh (**Fig. 5G-H**), yet show significant differential expression only in Punan Batu versus Punan Tubu and Lundayeh comparisons.

The state-transition dynamics of regulatory signals across the three contrasts further confirm this pattern. While strong combined signals of selection, expression, and methylation are concentrated in contrasts involving Punan Batu, the majority of regulatory states collapse to *none* in the Punan Tubu versus Lundayeh comparison (**Fig. S10**), consistent with the near-complete erasure of transcriptomic and methylome divergence between the transitioned hunter-gatherer and the agriculturalist. Fat-associated genes show a more pronounced collapse than non-fat-associated genes — 92% versus 86% transitioning to *none* (**Fig. S11**). Overall, this suggests lifestyle-driven expression plasticity following resettlement that is decoupled from the underlying selection signal and that genomic selection and transcriptional regulation are responding to largely distinct evolutionary pressures.

## Discussion

Human populations have shifted between foraging, mixed subsistence and agriculture over the last 10,000-15,000 years, yet the relative roles of genetic evolution, regulatory plasticity and epigenetic change in reconciling these transitions remain debated (Jeong and Di Rienzo, 2014). The Punan system provides an opportunity to disentangle the molecular consequences of rapid lifestyle change from deeper demographic history, because the Punan groups whom we work with in our study are closely related but differ in settlement and subsistence. We also work alongside an ancestrally distinct but ecologically similar agriculturalist neighbour, the Lundayeh. We are able to probe how quickly molecular systems respond to the lifestyle changes.

Phenotypic patterns across the hunter-gatherer — transitioned hunter-gatherer — agriculturalist gradient are consistent with a classic “nutrition and lifestyle transition” from mobile foraging to sedentary and market-integrated agriculture (Popkin, 1993, 2004). The relatively rapid emergence of fat-related phenotypic differences in the resettled Punan Tubu mirrors findings from other Indigenous populations in which short-term changes in diet, physical activity and infectious burden are linked to shifts in body composition and cardiometabolic risk (Gurven et al., 2013; Andersen et al., 2021; Gjermeni et al., 2025). These changes also fit within broader evolutionary models of metabolic imbalance, in which systems tuned to high-activity foraging ecologies become maladapted under conditions of energy surplus and reduced physical demand (Freese et al., 2018; Dounias and Froment, 2011). The enrichment of fat-related loci among genes showing strong differential expression indicates metabolism as one of the main axes of this transition even at the molecular level, consistent with published GWAS and eQTL studies implicating these genes in adiposity and cardiometabolic traits in other populations (Global Lipid Genetics Consortium, 2013; Locke et al., 2015; Shungin et al., 2015). Here, the most pronounced molecular markers of lifestyle transition may be transient and present in populations that reside in the intermediate positions along the forager-farmer spectrum.

Although the resettlement only happened 3-5 generations ago, transcriptomic and DNA methylation profiles of the Punan Tubu transitioned HG more closely resemble those of the Lundayeh agriculturalist than those of their Punan Batu hunter-gatherer relatives. This convergence indicates that gene expression and methylation states shift relatively rapidly with respect to lifestyle, manifesting molecular phenotypes that track current subsistence more than ancestry (Jaenisch and Bird, 2003; Colbran et al., 2021). Evolutionary modelling under an OU framework showed that the Punan Tubu have already moved approximately 77% of the way from the ancestral hunter-gatherer expression state toward the agriculturalist state. Furthermore, the dampening of molecular signals in the final post-transition state implies that once groups share a broadly similar lifestyle, molecular phenotype differences diminish, even where genomic ancestry remains distinct.

If transcriptional plasticity is sufficient to rapidly align the molecular phenotype of a transitioning population with that of established agriculturalists, this raises the question of whether positive selection on the underlying genomic sequence is necessary. Plastic responses that are repeatedly favoured by the environment may eventually be replaced or reinforced by genetic changes that produce the same phenotype (Morris, 2014; Coates et al., 2025), reducing the regulatory cost of maintaining plasticity (Pigliucci et al., 2006; Murren et al., 2015). Our data cannot resolve whether the Punan Tubu are on such a trajectory because the transition is too recent. However, the shared constraint strength *(α)* across populations in the OU model, combined with the limited selection-expression overlap, suggests that the regulatory architecture enabling this plastic response is not recently evolved. Plasticity, in this case, may be the primary and sufficient mechanism of short-term biological accommodation to lifestyle change, with genetic assimilation, if it occurs at all, operating on a much longer generational time.

Nevertheless, our study bears limitations that we acknowledge as follows. Our data are derived from whole blood which captures immune and metabolic signals robustly The metabolic and fat-related signals may reflect systemic regulatory changes in circulating immune cells responding to metabolic state and diets, rather than direct transcriptional changes in metabolically active tissues such as adipose, liver, or muscle. That said, blood gene expression has been shown to correlate with adiposity and metabolic traits (Glastonbury et al., 2016; Vôsa et al., 2021). Whether the fat-related gene expression shifts we observe in blood are mirrored in these metabolically active tissues remains an open question, and future studies incorporating tissuespecific profiling would strengthen the mechanistic interpretation of these findings. Secondly, the OU model assumes a star trifurcation with no post-split gene flow and a fixed shared root age. Although these assumptions are supported by our demographic analyses, deviations from them could change the estimated transition progress values. Lastly, while we interpret the rapid convergence as a reflection of plasticity, we cannot fully exclude the possibility that standing regulatory variation present in the ancestral Punan population prior to resettlement contributed to the speed of the observed shift. Longitudinal sampling of the transitioned groups across future generations would help distinguish ongoing plastic responses toward a new equilibrium.

In sum, our results show that the transition from hunting-gathering to agriculture in the Punan is accompanied by rapid directional changes in gene expression and DNA methylation, aligning the resettled Punan Tubu with long-term agriculturalists rather than their hunter-gatherer relatives. The speed and directionality of this shift highlight the importance of plasticity in mediating biological responses to lifestyle change. More broadly, our findings illustrate how recent lifestyle transitions can reshape human biology on timescales far shorter than those required for substantial genomic change, with direct implications for understanding and mitigating the health consequences of rapid economic and subsistence transitions in contemporary Indigenous populations worldwide.

## ACKNOWLEDGEMENTS

The authors thank the communities and volunteers involved in this research, as well as Datuk Abdul Karim, Antoni Ucan, and Sinang Luki for facilitating engagement with the communities in Punan Batu, Punan Tubu, and Lundayeh, respectively; Yayasan Konservasi Alam Nusantara; MRIN’s field team (Lidwina Priliani, Andreas Christian, and Isabella Apriyana), Kristiawan, and Kusaeri; and local health personnel in Puskesmas Bumi Rahayu, Puskesmas Malinau, and Puskesmas Pulau Sapi, Province of North Kalimantan, for the assistance during fieldwork and biological sampling. We also thank MRIN GDD lab members and SVI HGE lab members for comments and discussions. Funding was provided by the Wellcome Trust International Training Fellowship (no. 222992/Z/21/Z), Impact Seed Funding by the Pulitzer Center (to P.K.); the Leakey Foundation, the US National Science Foundation, and the Max Planck Institute for Evolutionary Anthropology (to J.S.L.); the National Health and Medical Research Council Ideas Grant 2020501 (to I.G.R.); the European Union’s Horizon 2020 research and innovation programme (MOBILE - no. 950610); and the UKRI ISPF ODA Grant (G128998 A33490) to G.S.J. St Vincent’s Institute acknowledges the infrastructure support it receives from the National Health and Medical Research Council Independent Research Institutes Infrastructure Support Program and from the Victorian Government through its Operational Infrastructure Support Program.

## CODE AND DATA AVAILABILITY

The whole genome sequence, whole transcriptome, and EPIC v2 DNA methylation data is currently being uploaded in European Genome-Phenome Archive (EGA). Raw data requests will be handled after formal publication with specific Data Access Agreement. Data S1-S4 are available at https://doi.org/10.5281/zenodo.21866451. The codes used for analysis are available at https://github.com/paikusuma/hg_transition.

## AUTHOR CONTRIBUTIONS

P.K., S.G.M., and I.G.R. conceived the study. P.K., G.S.J., F.A.M., P.C.L., and S.G.M. conducted the biological sampling. F.A.M. and P.C.L. provided project management. P.K. performed community engagement, laboratory work, bioinformatics data processing and statistical analyses, and wrote the initial manuscript draft. G.S.J. and H.L. advised on genetic analyses and their interpretations. J.S.L. advised on anthropological interpretations. I.G.R. advised on transcriptomic and DNA methylation data analysis and their interpretations. H.S. advised on the discussion. S.G.M. and I.G.R. provided supervision and advised on overall analyses, interpretations, and discussions. All authors contributed to the writing of the final manuscript.

## Methods

### Samples and Ethics

Here we report a new genomic dataset, which includes 78 high-coverage genomes from 3 populations in Indonesian Borneo, i.e. the Punan Batu Sajau community (n = 18) in Bulungan Regency, North Kalimantan Province; the Punan Tubu (n = 26) and Lundayeh (n = 34) communities in Malinau Regency, North Kalimantan Province. All samples were obtained from adult participants who agreed to participate in the research and filled out the informed consent. For full information about the new and published samples used in this study, refer to (**Table S1-S2**).

Community engagement was conducted with all three participating communities following principles of reciprocity and ongoing consent. Initial engagement involved dialogue with community elders and members to explain the aims of the research, discuss potential benefits and concerns, and clarify that participation was voluntary. Engagement was tailored to the specific context of each community. Across all three communities, free health screenings were provided in collaboration with local primary health clinics *(Puskesmas)*, offered to all community members regardless of their participation in the study to emphasise that the health check was an immediate benefit-sharing for the community and was non-coercive and independent of recruitment. Formal research permits were obtained from the relevant regional government authority (i.e., *Badan Kesbangpol* in Kabupaten Bulungan and Kabupaten Malinau, with support from the Department of Health from each Kabupaten via its Puskesmas) prior to fieldwork. Informed consent was obtained from all individual participants, with community-level consultation preceding individual consent processes.

The study as a whole, including community engagement, fieldwork, biological sample collection, and data analyses was reviewed and approved by the Mochtar Riady Institute for Nanotechnology Ethics Commission (Ethical Approval No. 006/MRIN-EC/ECL/III/2022, with Annual Continuing Review Approvals No. 012/MRIN-ECL/V/2023, and No. 011/MRIN-ECL/III/2024). Approval for data analysis in Australia was additionally granted by the University of Melbourne’s Human Research Ethics Committee (Approval ID 25179).

Blood samples were collected in EDTA vacutainers (for DNA) and Tempus Blood RNA Tubes (Applied Biosystems) (for RNA). DNA and RNA were extracted using Qiagen Puregene Kits and Tempus Spin RNA Isolation kit, respectively, following the manufacturer’s recommended protocol. RNA extractions were randomised with respect to populations. Quality and concentration of all extracted DNA samples were assessed using Qubit 4 (Life Technologies) and RNA samples using both TapeStation 4150 (Agilent) and Qubit 4 (Life Technologies). We only selected samples which had RIN (RNA Integrity Number) *>*5 for RNA sequencing.

### Phenotype Measurement

The research participants were measured for their weight using a body scale and their height using height rods. The waist and hip circumference were measured using a metric tape. Regarding the diet, the participants were interviewed using a modified semi-quantitative food frequency questionnaire. Blood glucose, triglyceride, total cholesterol (TC), and high- and low-density lipoprotein (HDL-LDL) were tested using point-of-care test strips.

For each individual, we recorded anthropometry (height, weight, BMI, waist and hip circumference) using height rods, a body scale, and a metric tape, respectively. We also recorded bioimpedance body-composition metrics (body fat, visceral fat, subcutaneous fat, muscle mass, and basal metabolic rate index) using Omron Karada Scan HBF-375 and blood biomarkers (glucose, total cholesterol, HDL, LDL, triglycerides, and uric acid) using point-of-care test strips.

Prior to analysis, continuous variables were inspected for outliers and distribution (using Shapiro-Wilk and Levene’s tests), log-transformed where appropriate to reduce skew, and standardised (mean = 0, SD = 1) for multivariate modelling. To identify phenotypes that differed significantly across lifestyle groups, we conducted Kruskal-Wallis tests for each variable, adjusting p-values for multiple comparisons using the Bonferroni method. For variables with overall p.adjust < 0.05, we performed post hoc pairwise Wilcoxon rank-sum tests with Bonferroni correction to determine which group contrasts (ternary: Punan Batu vs. Punan Tubu, Punan Batu vs. Lundayeh, Punan Tubu vs. Lundayeh; binary: HG (Punan Batu) vs. AGR (Punan Tubu and Lundayeh)) were significant. We computed nonparametric effect sizes to rank variables by magnitude of group difference. The top six variables by effect size from respective ternary and binary comparisons were retained for multivariate analyses.

To evaluate the capacity of the six phenotypes to predict lifestyle group, we implemented three supervised classifiers with 5-fold cross-validation using the `caret` package in R (Kuhn, 2008): i) Linear Discriminant Analysis (LDA): We trained an LDA model (method = “lda”) on the standardised predictors, using stratified 5-fold CV to estimate accuracy and Cohen’s *κ*, ii) Quadratic Discriminant Analysis (QDA): We trained a QDA model (method = “qda”) under the same CV scheme to allow group-specific covariance matrices. iii) Random Forest (RF): We trained an RF model (method = “rf”) with tuning of the `mtry` parameter (number of variables sampled at each split) over values around 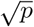, using 500 trees. We recorded out-of-fold accuracy, *κ*, and extracted permutation-based variable importance via `varImp()`. Model performance was compared by mean CV accuracy, *κ*, and class-specific sensitivity/specificity computed from the aggregated confusion matrix of out-of-fold predictions.

### Whole Genome

#### Sequencing, Base/Variant Calling, and Phasing

Sequencing libraries were prepared using Illumina DNA Prep PCR-Free. 150bp paired-end sequencing runs were performed on the Illumina NovaSeq X with an average of 600M reads per sample with the expected mean depth of 20x. The `fastp` (Chen, 2023; Chen et al., 2018) program was used to remove adapters, low-quality bases, and reads below 50bp, resulting in the minimum insert size of 350bp. The alignment was performed using `bwa-mem2` (Vasimuddin et al., 2019). We used the GRCh38 with alternative sequences, plus decoys and HLA for the alignment. All subsequent processes were either performed using `GATK v4.5` or Picard Tools within the `GATK v4.5` java framework (Auwera and O’Connor, 2020). After the alignment, fix mate information and mark duplicates was conducted using `FixMateInformation` and `MarkDuplicateSparks` commands, respectively. Following the MarkDuplicates step, the base quality scores were recalibrated and adjusted using `BaseRecalibrator` and `ApplyBQSR` commands. We used dbSNP 146 and Homo sapiens assembly 38 plus Mills and 1000G gold standard known indels for the base score recalibration, whose files can be found in the GATK resource bundle.

The base calling was undertaken using `GATK v4.5`’s `HaplotypeCaller`. The resulting gVCFs are consolidated into a GenomicsDB datastore. The `GenomicsDBImport` was performed in parallel on each chromosome. To capture worldwide genetic variants, the GenomicsDB datastore consists of this study’s newly generated data from Borneo and the published data from the Indonesian Genome Diversity Panel (Jacobs et al., 2019; Natri et al., 2022), 1000 Genomes (YRI, GBR, PJL, CHS, CDX, KHV, and PEL) (1000 Genomes Project Consortium et al., 2015), and Simon’s Genome Diversity Panel (Kankanaey, Ami, Dusun, and Papua New Guinea) (Mallick et al., 2016) **Table S1**, which included in the IGSR (Fairley et al., 2020).

Joint genotyping was performed using `GenotypeGVCFs`, outputting all sites to a multisample raw variant call. We then recalibrate the raw variants using `VariantRecalibrator` with hg38 hapmap 3.3, 1000genome, omni2.5, and high-confidence 1000 genome SNPs as training data for SNP, and Mills and 1000G gold standard known indels as training data for INDEL. We used a 99.7 truth-sensitivity filter for both SNP and INDEL and applied the filter using `ApplyVQSR`. Using `BCFtools v1.2`’s (Danecek et al., 2021) `setGT` plugins, the following filters were applied to each genotype call: base depth (DP) *<*8 and genotype quality (GQ) *<* 20 to set 0/0 genotypes into missing (· / ·). We then kept only biallelic variants. We masked sites within mappability (GRCh38 Mappability Stratification BED files) and low-complexity regions (GRCh38 Low-Complexity Stratification BED files). These mask files were downloaded from GIAB (https://github.com/genome-in-a-bottle/genome-stratifications/) (Olson et al., 2023). The genotypes were phased using `EAGLE v2.4.1` without reference (Loh et al., 2016). We added ancestral allele information to our dataset from the inferred EPO multiple alignments deposited in Ensembl’s 10 primates’ EPO (release 106).

#### Kinship and Population Structure Analyses

Kinship analysis was performed using `KING v.2.3.1` (Manichaikul et al., 2010). A portion of samples were excluded due to the presence of first-degree relatives in the dataset, indicated by the estimated kinship coefficient *>*0.17. Haplotype sharing using the `RefinedIBD` (Browning and Browning, 2013) was computed to estimate the total number of shared genetic fragments between each pair of individuals (LOD *>*3).

Principal Component Analysis (PCA) was performed, filtering out variants with minor allele frequency (MAF) of 0.05 and pruned for linkage disequilibrium with r^**2**^*>*0.2, using SNPRelate R package (Zheng et al., 2012). We used a range of D-statistics calculated using `ADMIXTOOLS v7.1` (Patterson et al., 2012) to interrogate the resettled Punan (Punan Tubu) and indigenous agricultural Borneans’ genetic affiliation to Punan Batu vs. the agricultural Lundayeh. D-statistics is a test of treeness based on allele frequency sharing between populations, on the form of *D(Outgroup, X* | *A, B)*. When the D values are significantly positive, it is indicative of gene flow occurred between the “*X*” and “*B*” group (relative to “*A*” group). In contrast, when negative, it is indicative of gene flow occurring between the “*X*” and “*A*” group (relative to “*B*” group). When the values are non-significant, the tree is considered balanced. We assigned African YRI as the outgroup and “*X*” as Punan Batu and relevant groups in the comparative dataset, while “*A*” and “*B*” are the Punan Batu and Lundayeh, respectively.

To estimate the population size dynamics, we conducted `MSMC2` analysis (Schiffels and Durbin, 2014). Based on `MSMC2`’s cross-coalescence results, we then performed the `MSMC-IM` (Wang et al., 2020) analyses to infer split times between a pair of populations. It fits a time-dependent migration model to the pairwise rate of coalescences based on estimates of coalescence rates within and across populations.

We also computed the cross-extended haplotype homozygosity (XP-EHH) (Sabeti et al., 2007) on the phased dataset using `selscan v.2.0.3` (Szpiech and Hernandez, 2014; Szpiech, 2024) to test for positive selection using haplotype information.

### Whole Transcriptome

#### Sequencing and Data Pre-processing

Sequencing libraries were prepared using TruSeq Stranded Total RNA with Ribo-Zero Globin. 150bp paired-end sequencing runs were performed on the Illumina NovaSeq X with an average of 60M paired reads per sample. Adapter removal and quality trimming were done using `Trim Galore` (Martin, 2011). Reads were aligned to the human genome (GRCh38.p14, GENCODE release 45) using `STAR v.27.11b` (Dobin et al., 2013) with two-pass alignment mode and GTF basic. There are more than 95% of uniquely mapped reads per sample. The Match Bam to VCF (MBV) method (Fort et al., 2017) from `QTLTools v.1.3` (Delaneau et al., 2017) was used to identify sample mix-ups and cross-contamination. MBV directly compares each aligned RNA-seq BAM file to all the genotypes in the VCF file and computes the proportion of concordant heterozygous and homozygous sites. Post-alignment QC was done using `RSeQC v5.0.1` (Wang et al., 2012). All reads have *>*92% paired-end antisense strands and *>*40% exonic reads. Read counts were then quantified with `featureCounts` (Liao et al., 2014) from `Subread` package release 2.0.6 (Liao et al., 2019) against a subset of GENCODE release 45’s GTF basic with transcript support levels 1-3. Read counts were converted to CPM (count per million) and TMM-normalised using `edgeR v4.2` (Robinson et al., 2010). Prior to TMM normalisation, we filtered out genes that have CPM *<*1 in 50% of each group, thus resulting in 13,710 genes.

#### Blood Cell Type Deconvolution

`Decon2`’s `DeconCell` (Aguirre-Gamboa et al., 2020) was used to estimate the proportion of CD8T, CD4T, NK, B cells, monocytes and granulocytes in each sample and tested these for association with the first 10 PCs of both the methylation and expression datasets. The normalisation and scaling for the deconvolution analysis used `DeconCell`’s `dCell.expProcessing()` function on the unnormalised CPM data. The proportion of each cell type was predicted using the normalised data and the reference bulk dataset.

#### Differential Expression

Linear models are fitted to the data with the assumption that the underlying data are normally distributed. A design matrix is set up with group/population, age, sex, RIN, rRNA ratio, and blood deconvolution proportion (i.e. CD8T, CD4T, NK, B cells, monocytes and granulocytes) information. Contrasts for 3 pairwise comparisons between populations (i.e.PunanBatu vs.PunanTubu; PunanBatu vs. Lundayeh; and PunanTubu vs. Lundayeh) are set up in `limma v3.60.6` (Ritchie et al., 2015) using the `makeContrasts()` function. We then removed heteroscedasticity from the count data using `limma`’s `voom()` (Law et al., 2014) function from the design matrix. Linear modelling is carried out using `limma`’s `lmFit()` and `contrasts.fit()` functions based on the design matrix and contrast matrix, respectively. Empirical Bayes moderation is then carried out following the linear modelling by borrowing information across all genes to obtain more precise estimates of gene-wise variability (Smyth, 2004).

Genes were called as differentially expressed (DEG) if the FDR-adjusted *p* value was below 0.05, regardless of the magnitude of the log_2_ fold change, unless noted otherwise. Lists of DEGs were annotated using `biomaRt v2.60.1` (Durinck et al., 2009). Gene set enrichment analyses for the DEGs were performed using `clusterProfiler v4.12` (Wu et al., 2021; Xu et al., 2024), with Gene Ontology and KEGG annotation drawn from the `org.Hs.eg.db v3.19` database.

#### Expression-Selection Gene Clustering and State-Transition

In order to classify genes according to their combined selection and expression profiles, we first constructed a six-dimensional feature matrix of -log_10_(*p*) values derived from XP-EHH and differential expression tests across three population contrasts (i.e., PunanBatu vs PunanTubu; PunanBatu vs Lundayeh; and PunanTubu vs Lundayeh). All six axes were standardised to zero mean and unit variance by *z*-score transformation. To identify the optimal number of clusters, we applied the elbow method on the total within-cluster sum of squares and computed average silhouette widths using the `silhouette()` function from the `cluster` package, followed by majority-rule consensus of 30 validity indices provided by the `NbClust` package (min.nc = 2, max.nc = 15). Genes were then partitioned using `stats::kmeans()` function with nstart = 25 random initialisations. Cluster quality was assessed via mean silhouette widths (values > 0 indicating well-separated clusters), and cluster centroids were inspected to define prototypical selection-expression trajectories.

For the state-transition analysis, we first encoded each gene’s status in each of the three population contrasts (Punan Batu vs Lundayeh, Punan Batu vs Punan Tubu, Punan Tubu vs Lundayeh) into 8 mutually exclusive states based on the presence or absence of three molecular signatures: evidence of positive selection *(sel)*, differential expression *(expr)*, differential methylation *(meth)*, and including all possible combinations of these features. Genes lacking any of these signatures were assigned to the *none* state. Genes classified as *none* in all three contrasts were excluded to focus on genes exhibiting at least one detectable signal. We summarised transition dynamics using several complementary metrics. For each phase, we calculated the proportion of genes assigned to non-*none* states, the fraction transitioning to *none*, and the proportion retaining the same non-*none* state across successive contrasts.

#### WGCNA

We performed weighted gene coexpression network analysis (WGCNA) (Langfelder and Horvath, 2008) to estimate correlations between suites of coexpressed genes and traits and groupings. WGCNA is an unbiased, data-driven method to cluster groups of genes with similar expression patterns. We used `limma`’s normalised counts above as the input for the WGCNA. We constructed a signed network with a soft thresholding power of 14 after `pickSoftThreshold()` function model with a scale-free topology fit (signed *R*^*2*^) > 0.8, and a minimum module size of 30 genes. We used dynamic tree cut and merged modules with greater than 75% similarity, producing a total of 19 modules. We plotted the FDR corrected Pearson correlation coefficient between module eigengenes and trait values.

#### Ornstein-Uhlenbeck Modelling of Evolutionary Dynamics

To characterise the evolutionary dynamics of lifestyle-associated gene expression differences, we fitted Ornstein-Uhlenbeck (OU) models to per-sample residualised expression values transformed with `voom`. Covariate effects (same variables as differential analyses) were removed by subtracting the fitted nuisance component from the `voom` log_2_-CPM matrix, yielding a residualised expression matrix retaining population signal and within-individual noise.

A star trifurcation topology was assumed, with all three populations diverging simultaneously from a common root. The root age was set to the mean of `MSMC-IM` pairwise lower-bound split times (PB-PT: 9,195 ya; PB-LDY: 10,677 ya; PT-LDY: 9,037 ya; mean: 9,636 ya; generation time: 29 years), giving *T*_*root*_ *=* 332 generations. No post-split gene flow was modelled, consistent with the absence of recent admixture between populations.

For each gene, five nested models were fitted by maximum likelihood. Under Brownian Motion (BM), expression evolves as a random walk with rate *σ*^*2*^, and all individuals share a common ancestral mean. Under the OU models, for each population *i*, expression is pulled toward a population-specific optimum *θ*_*i*_ at rate *α*, with tip variance:

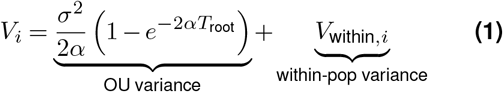

where *V*_*within,i*_ is the within-population individual variance. The five models were: BM (5 parameters: *σ*^2^, ancestral mean, *V*_*within*_ per population); OU with shared optimum and constraint (OU_shared_; 6 parameters: adds *α*); OU with population-specific optima (OU_*θ*_; 8 parameters: *θ*_*PB*_, *θ*_*PT*_, *θ*_*LDY*_ replace shared *θ*); OU with population-specific constraint strengths (OU_*α*_ ; 8 parameters: *α* _*PB*_, *α* _*PT*_, *α* _*LDY*_ replace shared *α*); and OU with both free (OU_*α θ*_; 10 parameters) (**Table S8**). Within-population variance *V*_*within,i*_ was estimated jointly with evolutionary parameters in all models rather than fixed from sample variance, allowing the likelihood to separate individual noise from between-population divergence.

All models were fitted by numerical optimisation using `optimx` in R, with both L-BFGS-B and Nelder-Mead methods and 20 random restarts per gene. Parameters *α*and *σ*^2^ were optimised on the log scale. Bounds on log *α* were set to [log(ln2*/*(100· *T*_*root*_)), 2], corresponding to half-lives between 0.09 and 33,170 generations, spanning the range from near-instantaneous to effectively neutral adaptation. Model selection used AIC, with the primary hypothesis test comparing OU*θ*against OU_shared_ via a likelihood ratio test (LRT; *χ*^*2*^, df = 2) with FDR correction. Models allowing population-specific *α* (OU_*α*_, OU_*α θ*_) were found to be unidentifiable across all genes (maximum likelihood gain *<* 10^−7^) and were excluded from downstream analyses.

For each gene, a transition progress metric was defined as *π =* (*θ*_*PT*_ − *θ*_*PB*_) / (*θ*_*LDY*_ − *θ*_*PB*_) for genes where | *θ*_*LDY*_ − *θ*_*PB*_ |*>* 0.1 log_2_-CPM, so that the ratio is computed over a meaningful dynamic range. Population-specific *θ* values were taken from the bestfitting model: OU_*θ*_where selected, and observed population means otherwise. The OU half-life was computed as *t*_*1/2*_ = ln2*/*_*α*_ in generations and converted to years using a generation time of 29 years. To assess whether PT’s expression shift toward LDY reflects lifestyle convergence independent of genetic background, we tested whether transition progress *π* predicts the probability that a gene differentially expressed between PB and LDY (FDR-adjusted *p <* 0.05) is no longer differentially expressed between PT and LDY, using logistic regression with *π* as the predictor.

### DNA Methylation

#### Array and Data Pre-processing

DNA methylation was conducted using Illumina Infinium MethylationEPIC v2.0 array scanned on Illumina iScan. Each methylation data point is represented by fluorescent signals from the M (methylated) and U (unmethylated) alleles. Background intensity computed from a set of negative controls was subtracted from each analytical data point. The ratio of fluorescent signals was then computed from the two alleles 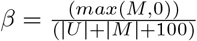 which reflects the methylation level of each CpG site. A *β*-value of 0 1 was reported signifying percent methylation, from 0% to 100%, respectively, for each CpG site.

Data *(β)* preprocessing was performed using `SeSAMe v1.22` (Ding et al., 2023; Zhou et al., 2018). The goal of this preprocessing step is to reduce systematic bias to avoid statistically erroneous conclusions through dye bias correction, low-quality probe masking, transformation, and *inferChannel* of data. We performed the recommended preprocessing for EPIC v2 using `openSesame()`’s prep code “QCDPB”, which does these functions in the following order: 1) masking (flagging) probes of poor design, 2) inferring channels for Infinium-I probes, 3) correcting dye bias using non-linear scaling, 4) masking p-value detection using oob (pOOBAH), and 5) subtracting background using oob. After that, we also excluded CpGs with at least one mask occurrence, excluded control probes & CpGs with NA values in at least one sample, and excluded probes on sex chromosomes. At first, there were 937,690 CpGs detected by the microarray. After the preprocessing, there were 880,992 CpGs.

#### Blood Cell Type Deconvolution

The deconvolution was initially performed using `SeSAMe`’s `estimateCellComposition()` function. The function takes a reference methylation status matrix (rows for probes and columns for cell types, can be obtained by `getRefSet()` function) and a query beta value measurement. However, the estimated results are unreliable as they deviate from the real biological phenomenon, e.g., the estimated CD19-B cells are close to zero for majority of the samples. Thus, we used instead RNA-seq’s blood deconvolution as covariates in downstream differential methylation analysis.

#### Differential Methylation

For differential analysis, we input M-values instead of *β* values, which translates 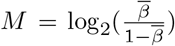. A design matrix is set up with group/population, age, sex, and blood deconvolution proportion derived from the expression data (i.e. CD8T, CD4T, NK, B cells, monocytes and granulocytes) information. Similar to the differential expression analysis, the differential methylation analysis was performed based on contrasts for 3 pairwise comparisons between populations (i.e., PunanBatu vs. PunanTubu; PunanBatu vs. Lundayeh; and PunanTubu vs. Lundayeh) using `limma v3.60.6` (Ritchie et al., 2015). Significant DMS (differentially methylated sites) were selected based on an FDR-adjusted *p* value threshold of 0.01 and a log_2_ fold change of 0.5 or greater. Enrichment tests for the DMS were performed using `missMethyl vl.38.0` (Phipson et al., 2016) `gometh()` function. Significantly enriched pathways were selected based on an FDR-adjusted *p*-value of 0.01.

### cis-eQTL and methylQTL mapping

SNPs with MAF *<* 0.05, call rate *<* 0.95 and Hardy-Weinberg equilibrium *p <* 0.0001 were removed in QTL mappings. As covariates, we included ten genotype principal components in QTL analyses to account for population structure. We used the probabilistic estimation of expression residuals (PEER) method (Stegle et al., 2010) to infer hidden sources of variation in expression and methylation data. These latent factors were used as surrogate variables for unknown technical batch effects and included as covariates in the QTL analyses. As many as 20 hidden factors (25% of the number of samples) were included in the models, as recommended in Stegle et al. (2012). Cis expression QTLs and cis methylation QTLs were called using `QTLTools v.1.3` (Delaneau et al., 2017). The covariates integrated in the regression model are listed and described in **Table S2**. Variants were defined as being in cis with a gene if they were located within a window of ±1Mb from the TSS or CpG for expression and methylation, respectively. We ran a permutation pass across chromosomes and performed FDR correction. We then ran subsequent conditional analysis from the nominal *p* thresholds (FDR-adjusted *p <* 0.05) obtained from the permutation pass.

