## Supplementary Figures and Tables for "Rapid convergence toward an agriculturalist regulatory landscape following lifestyle transition in Bornean hunter-gatherers"

### Supplementary Information

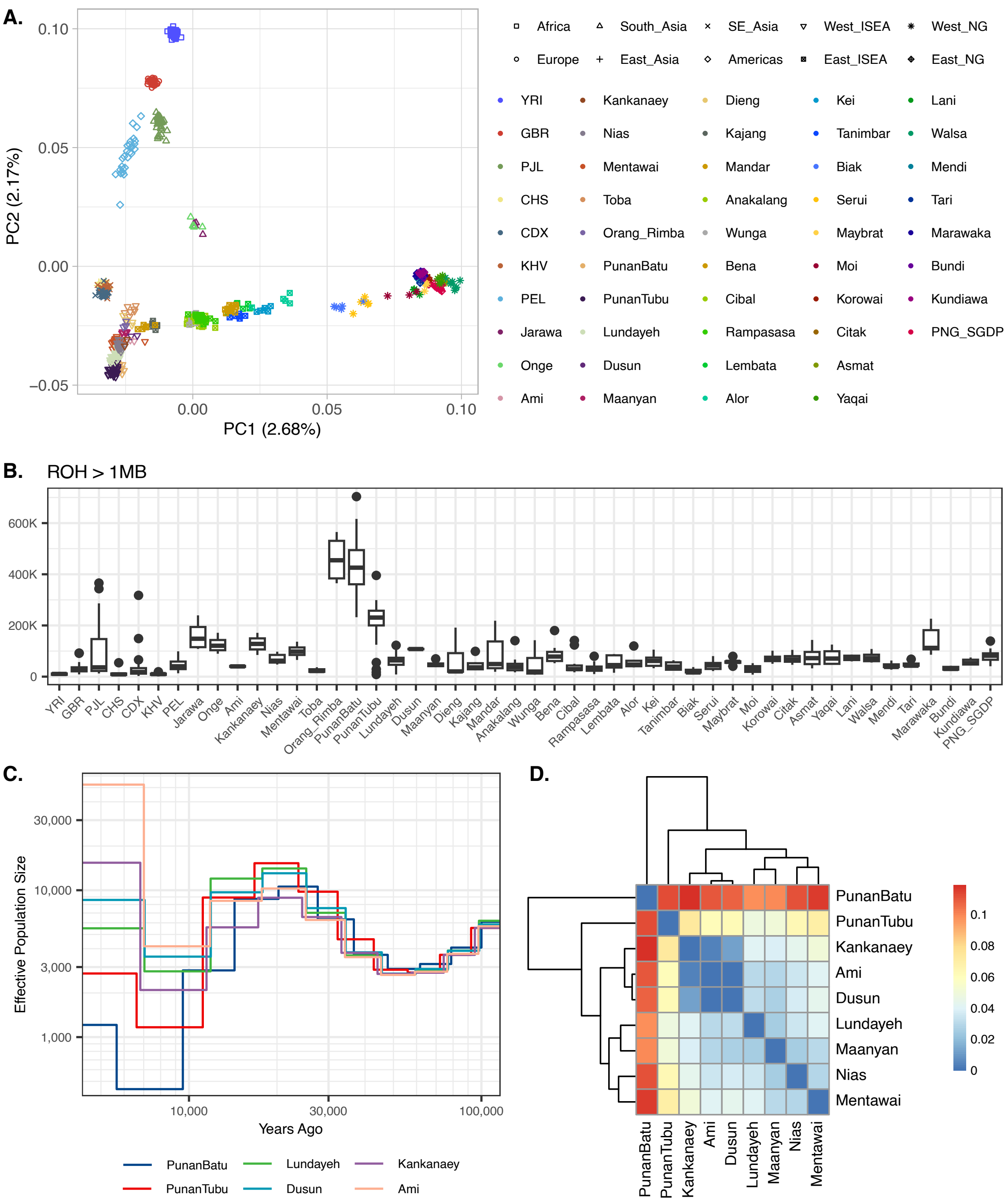


**Figure S1. Population structure, homozygosity, and demographics of Bornean populations** (**A**) PCA of genome data across all samples in the dataset, including Punan Batu (PB), Punan Tubu (PT), Lundayeh (LDY), and comparative populations. (**B**) Runs of Homozygosity (RoH) length distributions across populations, showing that Punan Batu carry substantially longer RoH segments than other Bornean groups and at comparable levels to a hunter-gatherer population (Orang Rimba) in the region. (**C**) Dynamics of effective population size ($N_e$) through time estimated from MSMC2, showing the demographic trajectories of PB, PT, and LDY alongside comparative populations. (**D**) Pairwise $F_{ST}$ values across population comparisons.

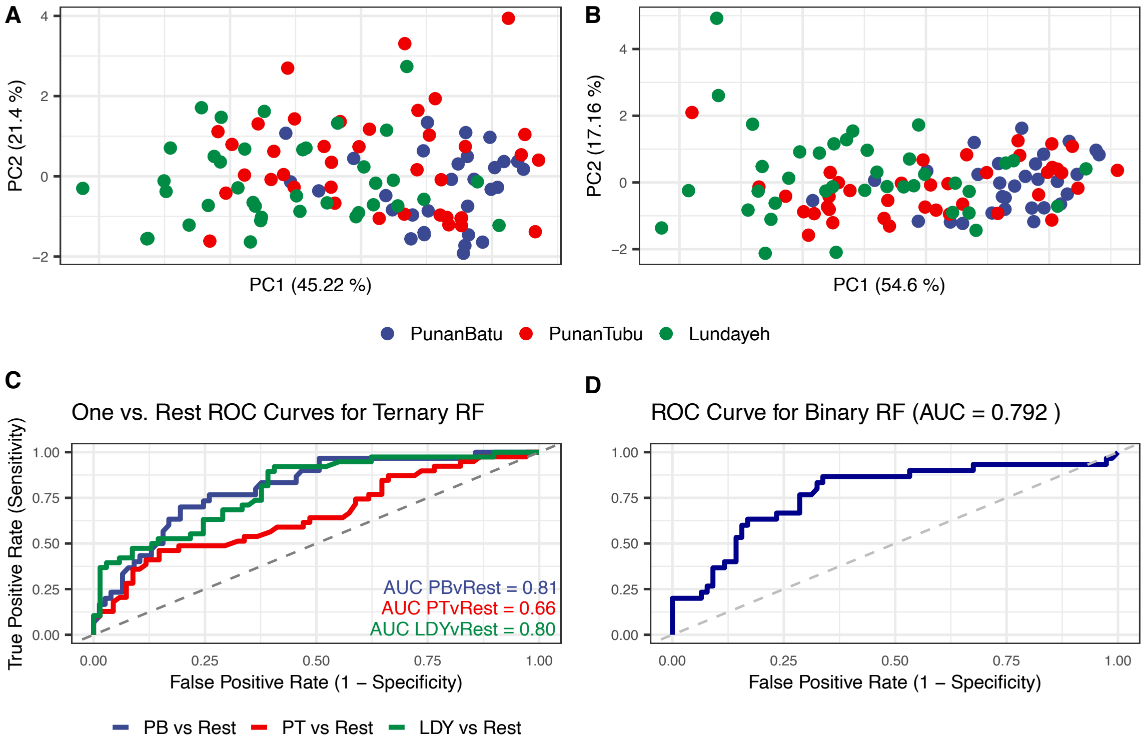


**Figure S2. Random Forest classifier analysis for phenotype data for terniary and binary groups**

(**A**) PCA of the top phenotypes values for the ternary classifier across all three classes. (**B**) PCA of the top phenotypes values for the binary hunter-gatherer versus transitioned classifier. (**C**) ROC curves for the ternary classifier across all three classes under 5-fold cross-validation. (**D**) ROC curves for the binary hunter-gatherer versus transitioned classifier under 5-fold cross-validation.

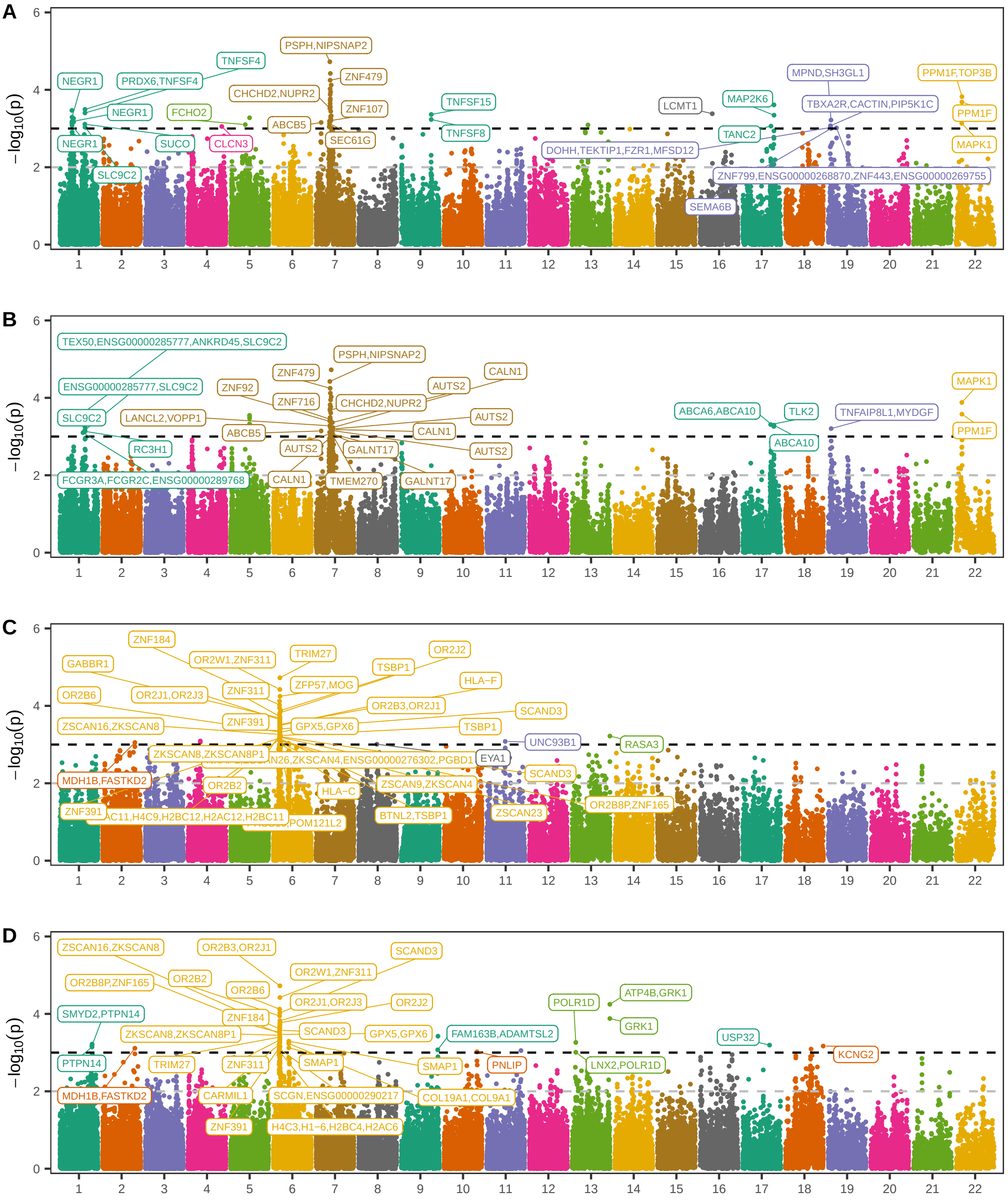


**Figure S3. Positive selection scans reveal population-specific signatures largely independent of lifestyle transition.**
(**A–B**) XP-EHH selection scans comparing Punan Batu against Lundayeh (**A**) and Punan Tubu (**B**), with consistent peaks on chromosome 7 harbouring *PSPH* and chromosome 22 harbouring *MAPK1* appearing in both contrasts. (**C–D**) XP-EHH selection scans comparing Punan Tubu against Punan Batu (**C**) and Lundayeh (**D**), with a consistent peak on chromosome 6 near a cluster of olfactory receptor genes (*OR2* family) appearing in both contrasts.

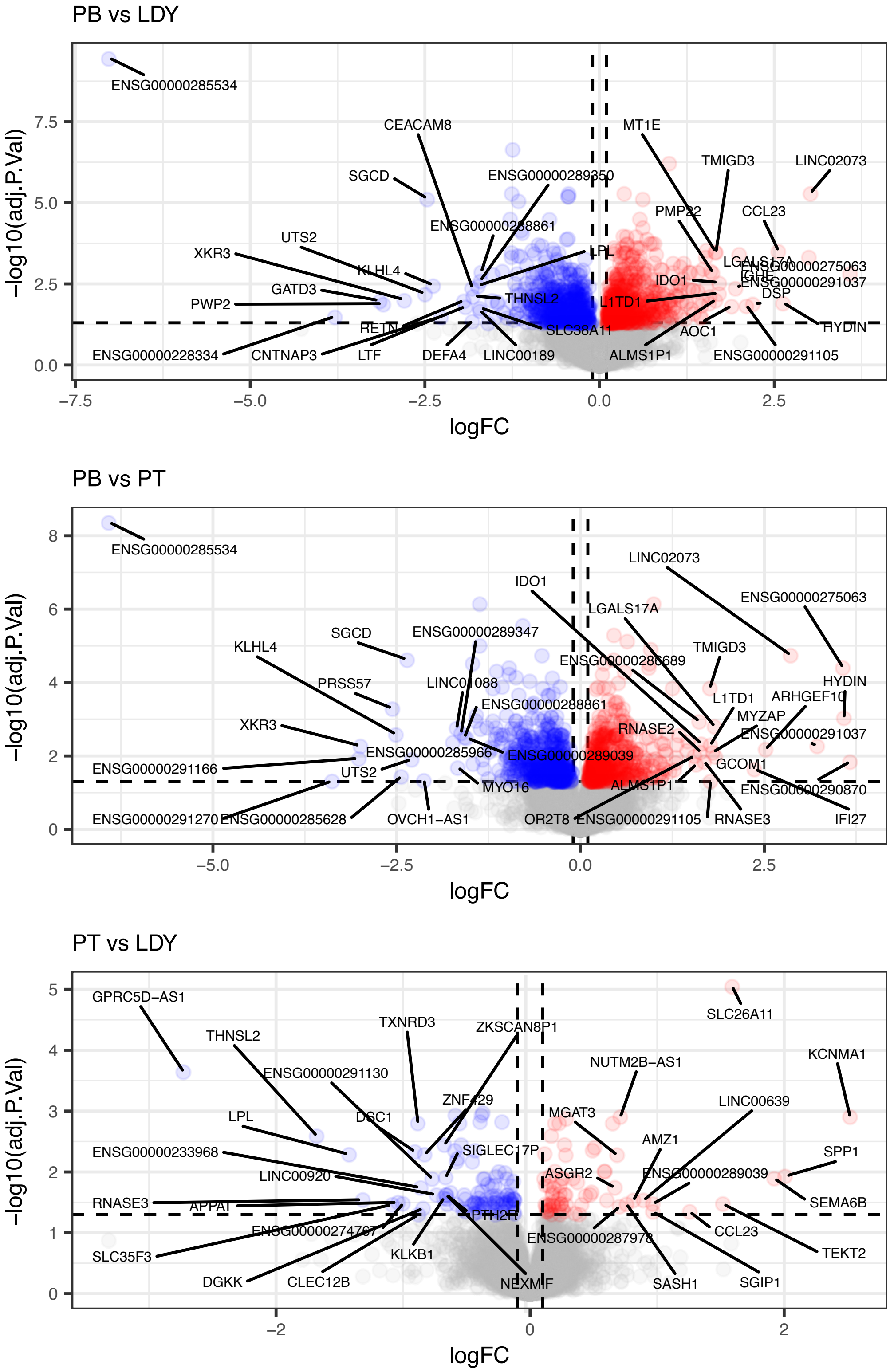


**Figure S4. Differential expression landscape across pairwise population contrasts.** (**A–C**) Volcano plots of differential gene expression for Punan Batu versus Lundayeh (**A**), Punan Batu versus Punan Tubu (**B**), and Punan Tubu versus Lundayeh (**C**), showing $-\log_{10}$ FDR-adjusted $p$-values against $\log_2$ fold-change for all expressed genes. Significantly differentially expressed genes (FDR-adjusted $p <$ 0.05, fold-change $\geq$ 1.1) are highlighted, with genes upregulated and downregulated relative to the first-named population shown in distinct colours.

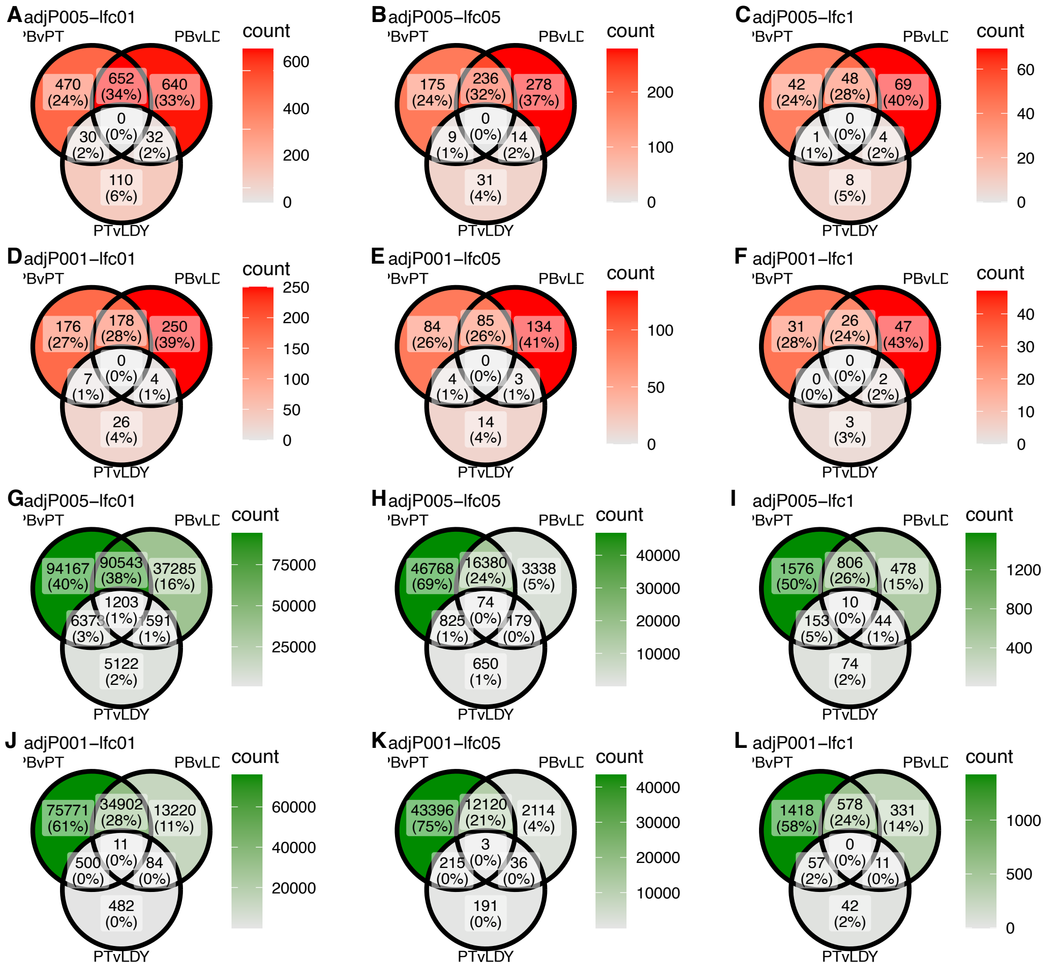


**Figure S5. Overlap of differentially expressed genes and differentially methylated sites across pairwise contrasts at varying significance and effect size thresholds.**
(**A–F**) Venn diagrams of differentially expressed genes (DEGs) across the three pairwise contrasts — Punan Batu versus Lundayeh (PB vs LDY), Punan Batu versus Punan Tubu (PB vs PT), and Punan Tubu versus Lundayeh (PT vs LDY) — at FDR-adjusted $p < 0.05$ with $\log_2$FC thresholds of 0.1 (**A**), 0.5 (**B**), and 1.0 (**C**), and at FDR-adjusted $p < 0.01$ with the same $\log_2$FC thresholds of 0.1 (**D**), 0.5 (**E**), and 1.0 (**F**). (**G–L**) Venn diagrams of differentially methylated sites (DMS) across the same three pairwise contrasts at FDR-adjusted $p < 0.05$ with $\log_2$FC thresholds of 0.1 (**G**), 0.5 (**H**), and 1.0 (**I**), and at FDR-adjusted $p < 0.01$ with thresholds of 0.1 (**J**), 0.5 (**K**), and 1.0 (**L**). Across all thresholds and both molecular layers, the overlap across all three contrasts is consistently minimal, and the PT versus LDY comparison contributes the fewest unique and shared signals, confirming that the depletion of differential signals between the two post-transition groups is a robust rather than an artefact of any particular significance or effect size threshold.

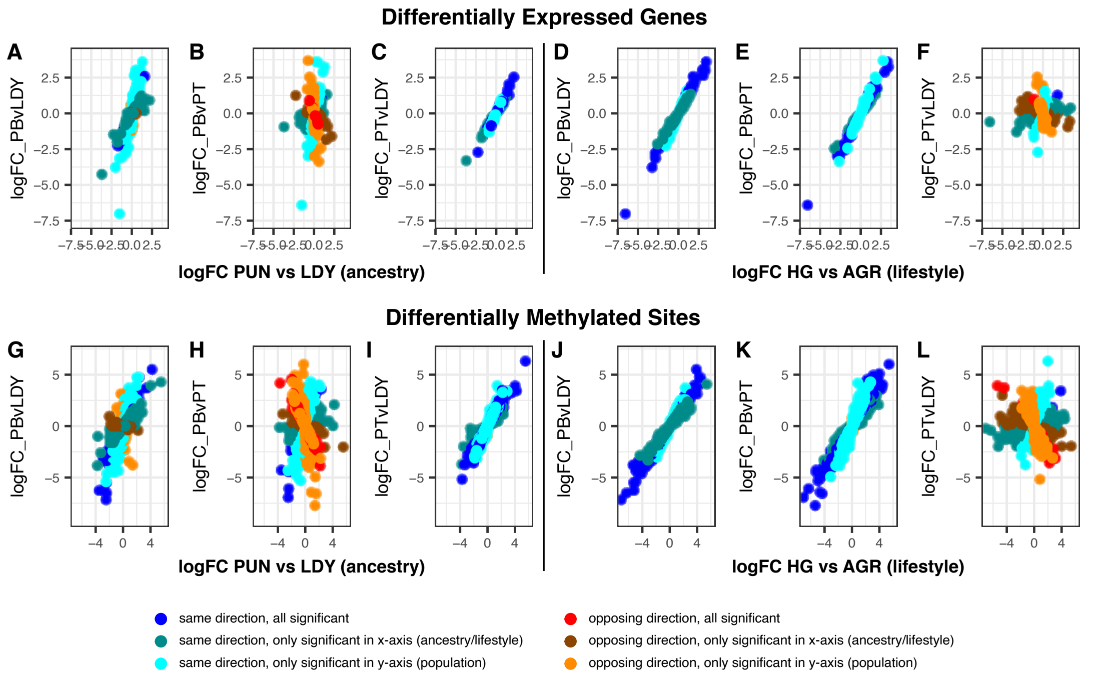


**Figure S6. Lifestyle rather than ancestry drives the directional concordance of transcriptomic and epigenetic divergence.**

Scatter plots comparing $\log$FC values across ancestry-based and lifestyle-based stratifications for differentially expressed genes (DEGs; **A–F**) and differentially methylated sites (DMS; **G–L**). Left panels (**A–C**, **G–I**) show ancestry-based comparisons, where the x-axis represents the combined Punan groups versus Lundayeh (HG/TransHG vs Ag) and y-axes show pairwise $\log$FC values for PB vs LDY, PB vs PT, and PT vs LDY respectively. Right panels (**D–F**, **J–L**) show lifestyle-based comparisons, where the x-axis represents Punan Batu versus the combined post-transition groups (HG vs TransHG/Ag) and y-axes show the same pairwise $\log$FC values. Genes and sites showing concordant directionality — where the sign of $\log$FC is consistent across comparisons — are more prevalent in the lifestyle stratification than in the ancestry stratification, indicating that lifestyle transition is the dominant axis of molecular divergence in this system. Discordant signals in the ancestry-based panels, particularly in the PB vs LDY versus PT vs LDY comparison (**A**, **G**), identify loci where Punan Tubu has shifted toward the Lundayeh profile relative to Punan Batu.

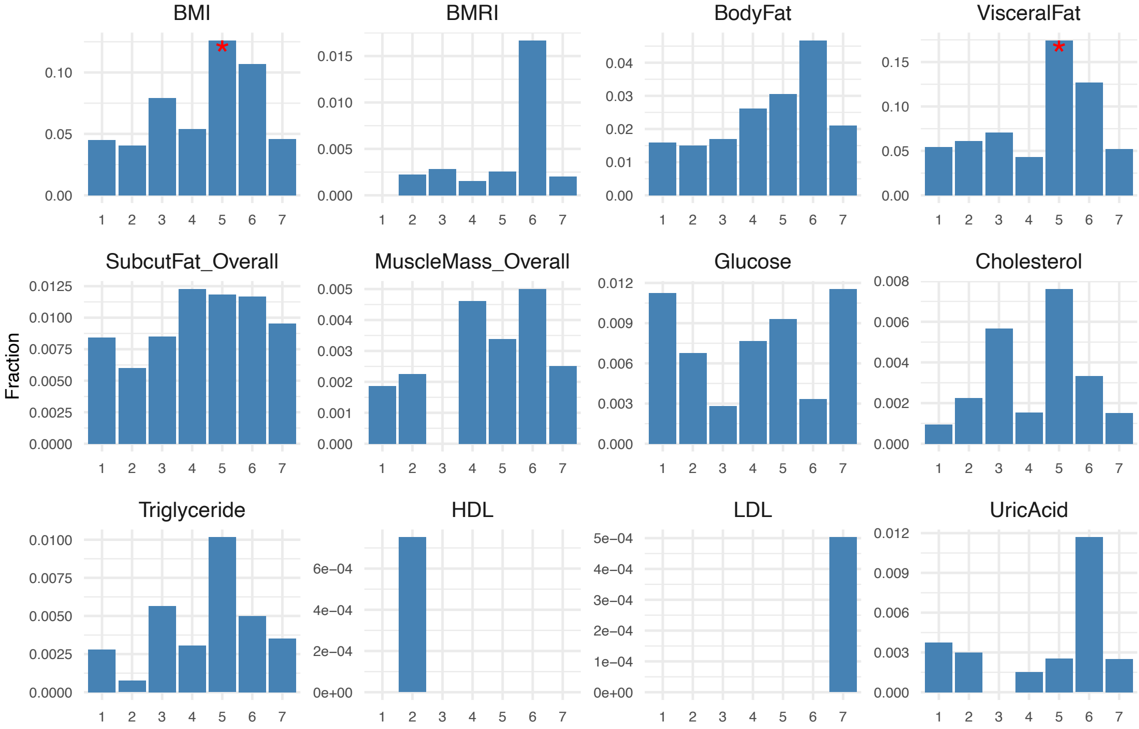


**Figure S7. Phenotype enrichment of $k$-means clusters identifies fat metabolism genes as the primary molecular signature of lifestyle transition.**
Fraction of phenotype-associated genes in each of the seven $k$-means clusters (x-axis) for twelve traits: BMI, basal metabolic rate index (BMRI), body fat, visceral fat, subcutaneous fat, muscle mass, blood glucose, cholesterol, triglycerides, HDL, LDL, and uric acid. Each panel shows the proportion of genes in a given cluster that are associated with the corresponding trait, with asterisks indicating significant enrichment relative to the genome-wide background (Fisher's exact test, FDR-adjusted $p < 0.05$).

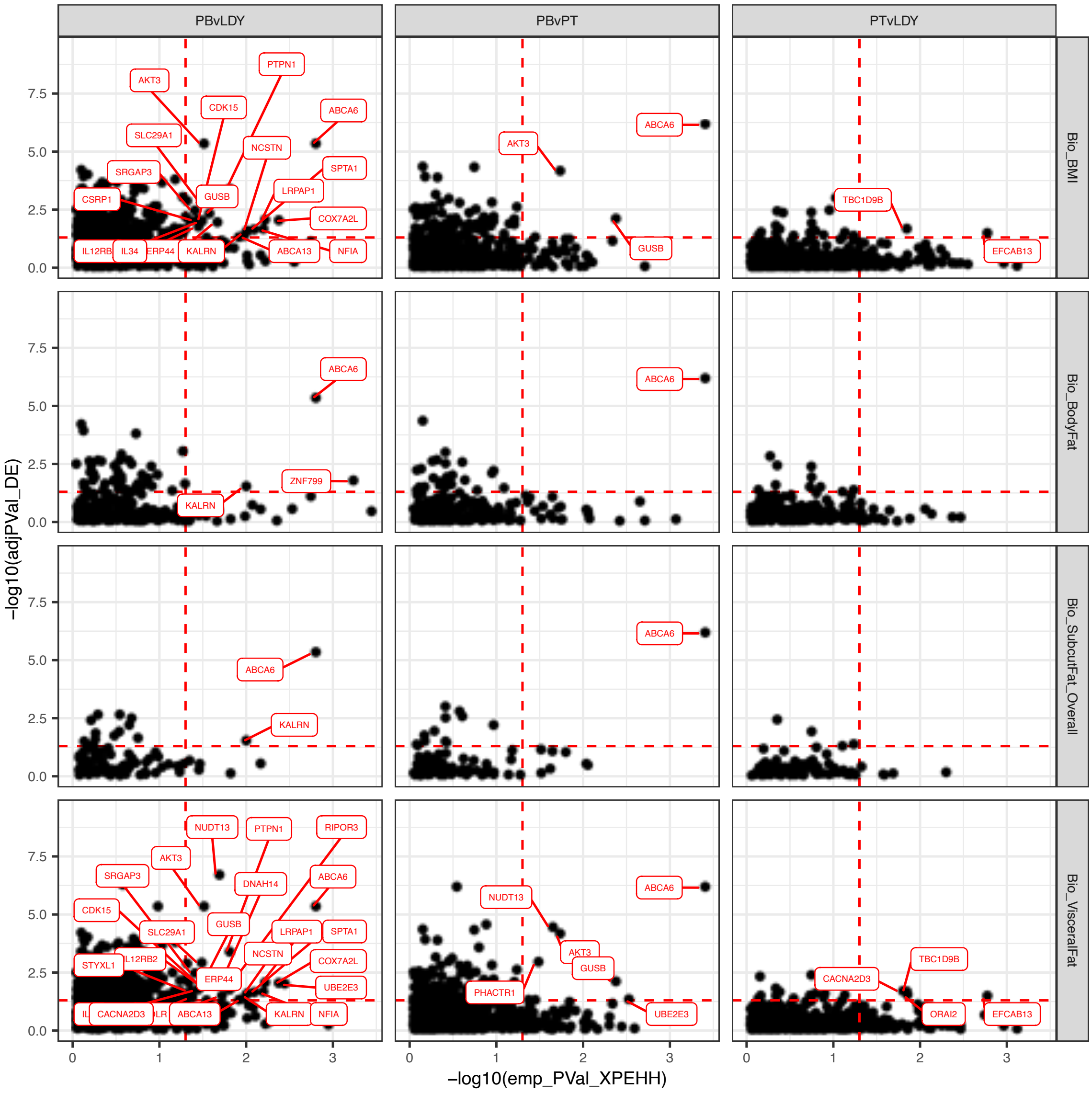


**Figure S8. Integration of selection signatures and phenotype-associated gene expression.** Faceted scatter plots integrating XP-EHH selection signals with differential gene expression associated with metabolic phenotypes. Each point represents a gene. The x-axis shows $-\log_{10}$ empirical p-values from cross-population extended haplotype homozygosity (XP-EHH) scans, indicating strength of positive selection. The y-axis shows $-\log_{10}$ adjusted $p$-values for differential gene expression (DEGs). Vertical facets represent population contrasts: Punan Batu (HG) vs Lundayeh (Ag), Punan Batu vs Punan Tubu (TransHG), and Punan Tubu vs Lundayeh. Horizontal facets represent phenotypic categories: body mass index (BMI), body fat percentage, subcutaneous fat, and visceral fat. Genes positioned in the upper-right quadrant show both strong selection signals and significant differential expression associated with metabolic phenotypes.

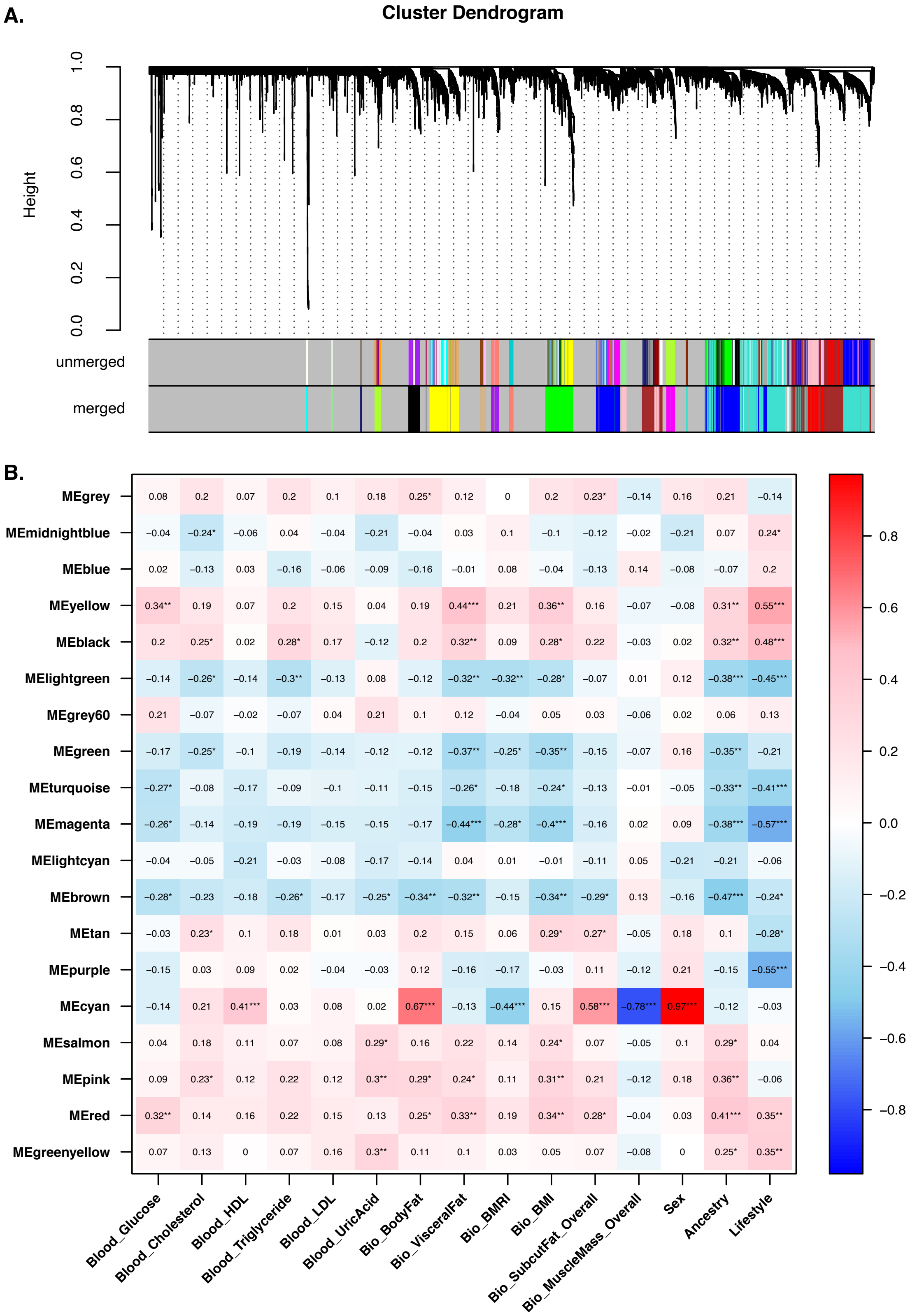


**Figure S9. Co-expression network analysis identifies coordinated gene modules associated with lifestyle, ancestry, and adiposity traits.**

(**A**) Cluster dendrogram of all expressed genes constructed by Weighted Gene Co-expression Network Analysis (WGCNA), with module colour assignments indicating groups of co-expressed genes. (**B**) Module-trait correlation heatmap showing Pearson correlations between module eigengenes and phenotypic and population-level traits.

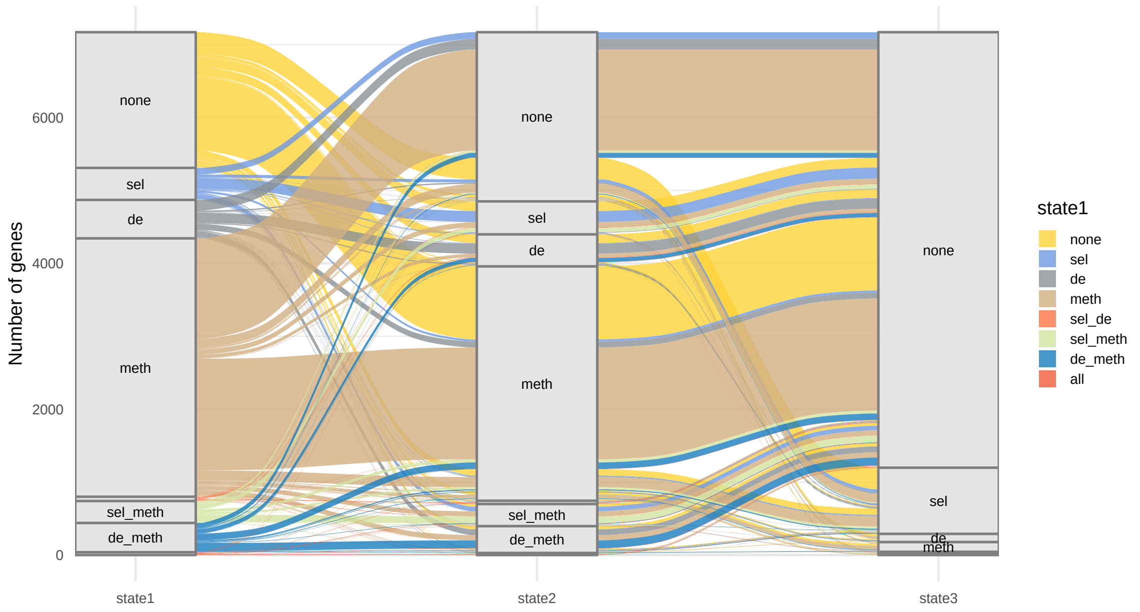


**Figure S10.** Alluvial plot showing transitions of gene states across three pairwise contrasts: ancestral HG state (PBvLDY), mid-transition (PBvPT), and post-transition (PTvLDY). Each stratum represents one of 8 possible states defined by combinations of selection (*sel*), differential expression (*expr*), differential methylation (*meth*), and their intersections. Flows represent the number of genes transitioning between states across contrasts. Genes assigned as *none* in all three contrasts were excluded.

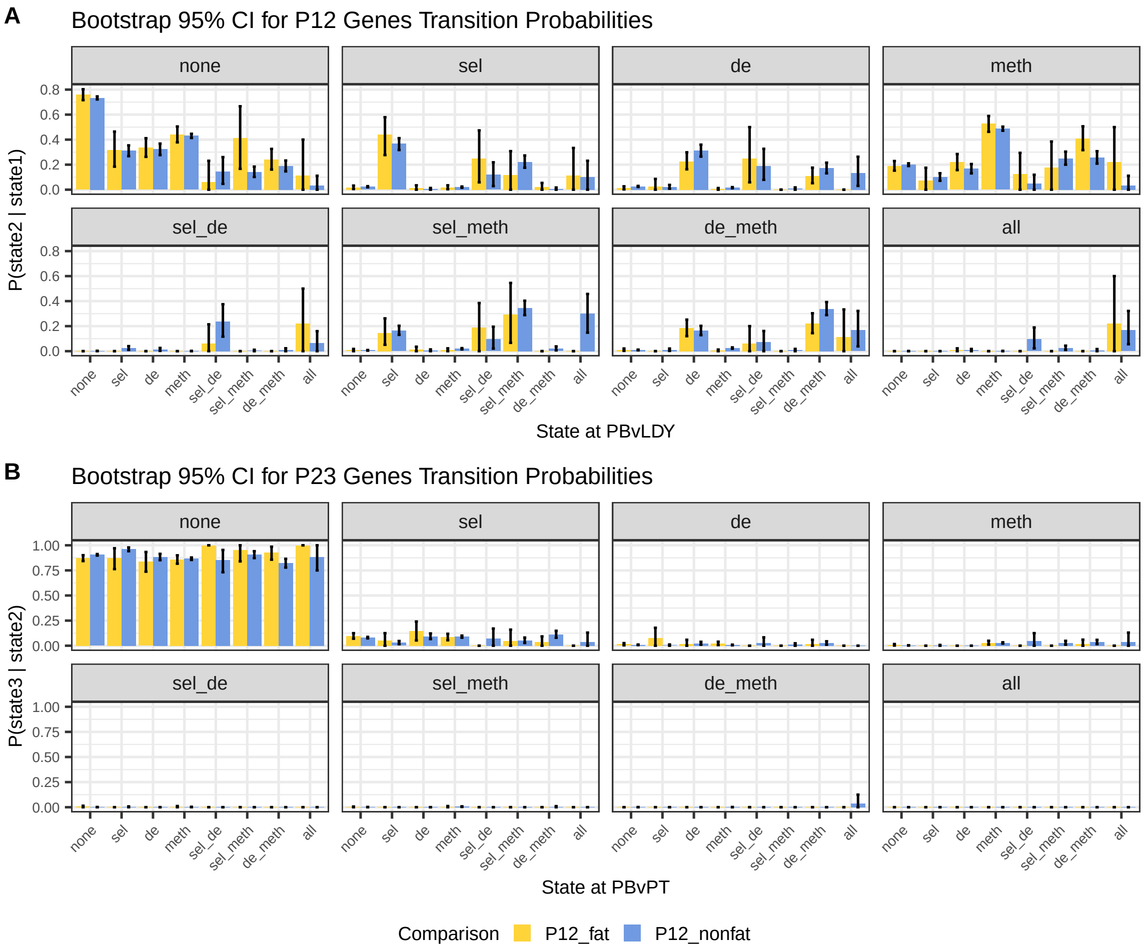


**Figure S11. Bootstrap-derived transition probabilities reveal asymmetric regulatory dynamics across lifestyle contrasts.**

(**A**) Bootstrap 95% confidence intervals for all gene state transition probabilities from State 1 (PB vs LDY) to State 2 (PB vs PT), shown separately for fat-associated (orange) and non-fat-associated (blue) genes across all 8 states defined by combinations of positive selection (*sel*), differential expression (*expr*), methylation (*meth*), and their intersections. Points represent estimated transition proportions, and error bars represent bootstrap-derived 95% confidence intervals from 1,000 resampling iterations. (**B**) Bootstrap 95% confidence intervals for gene state transition probabilities from State 2 (PB vs PT) to State 3 (PT vs LDY), shown for the same gene categories and state definitions. The State 2 to State 3 transition is dominated by convergence toward the *none* state across both gene categories.

### Supplementary Tables

| No | Population | Region | Dataset | n() |
|---|---|---|---|---|
| 1 | YRI | Africa | 1000 Genomes | 90 |
| 2 | PEL | Americas | 1000 Genomes | 90 |
| 3 | CHS | East_Asia | 1000 Genomes | 90 |
| 4 | GBR | Europe | 1000 Genomes | 90 |
| 5 | CDX | SE_Asia | 1000 Genomes | 90 |
| 6 | KHV | SE_Asia | 1000 Genomes | 90 |
| 7 | PJL | South_Asia | 1000 Genomes | 90 |
| 8 | Jarawa | South_Asia | Mondal et al. 2016 | 4 |
| 9 | Onge | South_Asia | Mondal et al. 2016 | 6 |
| 10 | Lundayeh | West_ISEA | this study | 36 |
| 11 | PunanBatu | West_ISEA | this study | 26 |
| 12 | PunanTubu | West_ISEA | this study | 34 |
| 13 | Ami | West_ISEA | SGDP | 2 |
| 14 | Kankanaey | West_ISEA | SGDP | 2 |
| 15 | Dusun | West_ISEA | SGDP | 2 |
| 16 | Toba | West_ISEA | Jacobs et al. 2019 | 7 |
| 17 | Mentawai | West_ISEA | Jacobs et al. 2019 | 39 |
| 18 | Nias | West_ISEA | Jacobs et al. 2019 | 16 |
| 19 | OrangRimba | West_ISEA | Jacobs et al. 2019 | 4 |
| 20 | Dieng | West_ISEA | Jacobs et al. 2019 | 7 |
| 21 | Maanyan | West_ISEA | Jacobs et al. 2019 | 7 |
| 22 | Kajang | East_ISEA | Jacobs et al. 2019 | 6 |
| 23 | Mandar | East_ISEA | Jacobs et al. 2019 | 6 |
| 24 | Alor | East_ISEA | Jacobs et al. 2019 | 5 |
| 25 | Cibal | East_ISEA | Jacobs et al. 2019 | 1 |
| 26 | Lembata | East_ISEA | Jacobs et al. 2019 | 7 |
| 27 | Bena | East_ISEA | Jacobs et al. 2019 | 12 |
| 28 | Cibal | East_ISEA | Jacobs et al. 2019 | 12 |
| 29 | Rampasasa | East_ISEA | Jacobs et al. 2019 | 19 |
| 30 | Anakalang | East_ISEA | Jacobs et al. 2019 | 15 |
| 31 | Wunga | East_ISEA | Jacobs et al. 2019 | 15 |
| 32 | Tanimbar | East_ISEA | Jacobs et al. 2019 | 6 |
| 33 | Kei | East_ISEA | Jacobs et al. 2019 | 6 |
| 34 | Korowai | West_NG | Jacobs et al. 2019 | 7 |
| 35 | Lani | West_NG | this study | 4 |
| 36 | Citak | West_NG | Jacobs et al. 2019 | 4 |
| 37 | Korowai | West_NG | Jacobs et al. 2019 | 7 |
| 38 | PNG_SGDP | East_NG | SGDP | 12 |
| 39 | Bundi | East_NG | Malaspinas et al. 2016 | 5 |
| 40 | Kundiawa | East_NG | Malaspinas et al. 2016 | 5 |
| 41 | Marawaka | East_NG | Malaspinas et al. 2016 | 5 |
| 42 | Mendi | East_NG | Malaspinas et al. 2016 | 5 |
| 43 | Tari | East_NG | Malaspinas et al. 2016 | 5 |

**Table S1. List of populations involved in the analyses of this study.**

| SampleID | Group | Extr Batch | RIN | rRNA ratio | Age | Sex | Lane | Granulo | B CD19+ | T CD4+ | T CD8 | NK CD3-CD56+ | Mono CD14+ |
|---|---|---|---|---|---|---|---|---|---|---|---|---|---|
| BLGII_003 | PunanBatu | 3 | 5.2 | 0.7 | 60 | M | 2 | 45.93 | 3.57 | 21.1 | 10.32 | 6.36 | 6.82 |
| BLGII_004 | PunanBatu | 4 | 4.9 | 1.6 | 32 | M | 1 | 48.89 | 2.92 | 20.91 | 9.26 | 5.04 | 6.58 |
| BLGII_005 | PunanBatu | 1 | 4.9 | 0.5 | 38 | M | 2 | 49.50 | 3.46 | 22.22 | 8.80 | 4.56 | 6.53 |
| BLGII_008 | PunanBatu | 1 | 5.2 | 2.2 | 60 | F | 1 | 48.08 | 3.51 | 22.41 | 8.77 | 4.20 | 6.61 |
| BLGII_009 | PunanBatu | 2 | 6.2 | 1.7 | 65 | F | 2 | 49.08 | 3.29 | 22.49 | 9.21 | 3.54 | 6.56 |
| BLGII_010 | PunanBatu | 6 | 4.9 | 0.9 | 35 | F | 1 | 51.21 | 2.05 | 17.61 | 10.40 | 3.55 | 7.36 |
| BLGII_014 | PunanBatu | 3 | 4.9 | 0.4 | 55 | F | 2 | 45.47 | 3.57 | 22.44 | 10.49 | 5.71 | 6.31 |
| BLGII_015 | PunanBatu | 5 | 5.7 | 1.1 | 40 | M | 2 | 53.18 | 2.93 | 20.93 | 11.99 | 5.40 | 6.14 |
| BLGII_017 | PunanBatu | 7 | 6.7 | 3 | 25 | F | 1 | 52.05 | 2.63 | 19.45 | 11.01 | 4.02 | 6.74 |
| BLGII_018 | PunanBatu | 1 | 5.7 | 1 | 20 | F | 2 | 51.93 | 2.33 | 18.65 | 8.13 | 3.52 | 7.45 |
| BLGII_019 | PunanBatu | 2 | 6.3 | 1.5 | 70 | M | 1 | 51.68 | 2.65 | 17.57 | 8.50 | 5.75 | 7.62 |
| BLGII_020 | PunanBatu | 3 | 5.5 | 0.9 | 40 | M | 1 | 49.41 | 3.28 | 20.44 | 9.32 | 4.58 | 6.67 |
| BLGII_021 | PunanBatu | 7 | 8.2 | 1.5 | 22 | M | 2 | 53.34 | 3.03 | 20.75 | 8.82 | 4.11 | 6.52 |
| BLGII_022 | PunanBatu | 4 | 5.8 | 0.8 | 48 | F | 1 | 53.95 | 2.67 | 19.92 | 8.41 | 4.37 | 7.05 |
| BLGII_023 | PunanBatu | 5 | 5.6 | 1.4 | 40 | F | 1 | 46.37 | 2.86 | 23.02 | 10.37 | 5.46 | 6.15 |
| BLGII_027 | PunanBatu | 6 | 4.9 | 1.9 | 43 | M | 2 | 50.43 | 3.07 | 21.99 | 9.20 | 5.40 | 6.44 |
| BLGII_028 | PunanBatu | 4 | 5.7 | 2.2 | 65 | M | 1 | 47 | 3.27 | 21.32 | 10.10 | 5.40 | 6.52 |
| BLGII_029 | PunanBatu | 3 | 5.8 | 1.3 | 20 | M | 1 | 51.14 | 2.69 | 20.06 | 11.33 | 4.60 | 6.18 |
| MAL_RPNII_001 | PunanTubu | 4 | 5.7 | 1.3 | 60 | M | 2 | 52.71 | 2.49 | 19.32 | 8.01 | 6.30 | 6.99 |
| MAL_RPNII_002 | PunanTubu | 1 | 6 | 1.2 | 51 | F | 2 | 47.51 | 2.69 | 22.06 | 10.42 | 5.35 | 6.80 |
| MAL_RPNII_004 | PunanTubu | 5 | 7.3 | 2.1 | 49 | M | 1 | 54.42 | 2.36 | 19.85 | 9.07 | 4.71 | 6.87 |
| MAL_RPNII_005 | PunanTubu | 3 | 6.5 | 2.3 | 48 | F | 1 | 54.17 | 1.86 | 18.48 | 9.29 | 5.27 | 6.79 |
| MAL_RPNII_006 | PunanTubu | 2 | 5.9 | 1.2 | 70 | F | 2 | 48.48 | 2.61 | 18.61 | 8.96 | 5.85 | 6.80 |
| MAL_RPNII_007 | PunanTubu | 5 | 5.5 | 1.3 | 63 | F | 1 | 48.84 | 2.73 | 19.35 | 9.04 | 6.90 | 6.43 |
| MAL_RPNII_008 | PunanTubu | 4 | 7.8 | 1.5 | 34 | M | 2 | 52.24 | 1.44 | 17.89 | 8.73 | 5.36 | 7.05 |
| MAL_RPNII_009 | PunanTubu | 5 | 7.6 | 2.2 | 52 | M | 1 | 46.51 | 2 | 21.37 | 9 | 6.07 | 6.77 |
| MAL_RPNII_010 | PunanTubu | 1 | 6.4 | 1.3 | 56 | M | 2 | 55.54 | 2.28 | 18.48 | 8.49 | 2.49 | 7.43 |
| MAL_RPNII_011 | PunanTubu | 1 | 7.3 | 1.7 | 55 | M | 1 | 59.07 | 1.85 | 16.05 | 7.90 | 5.18 | 7.39 |
| MAL_RPNII_016 | PunanTubu | 3 | 5.6 | 1 | 71 | F | 1 | 48.80 | 2.18 | 21.46 | 9.66 | 4.37 | 6.91 |
| MAL_RPNII_017 | PunanTubu | 7 | 7 | 1.4 | 47 | F | 1 | 52.98 | 2.26 | 21 | 10.93 | 3.86 | 6.59 |
| MAL_RPNII_019 | PunanTubu | 4 | 5.8 | 1.4 | 19 | F | 1 | 51.67 | 2.72 | 19.66 | 9.44 | 3.83 | 6.84 |
| MAL_RPNII_020 | PunanTubu | 4 | 5.9 | 1.4 | 59 | F | 1 | 47.30 | 2.99 | 23.73 | 9.18 | 4.81 | 6.14 |
| MAL_RPNII_021 | PunanTubu | 3 | 5.5 | 2 | 36 | F | 2 | 51.24 | 2.13 | 19 | 9.06 | 6.29 | 7.08 |
| MAL_RPNII_025 | PunanTubu | 1 | 8.1 | 2 | 47 | M | 1 | 58.39 | 2.25 | 18.83 | 7.64 | 4.20 | 6.95 |
| MAL_RPNII_028 | PunanTubu | 4 | 5.6 | 1.3 | 59 | F | 1 | 49.27 | 2.96 | 21.13 | 8.72 | 5.49 | 6.49 |
| MAL_RPNII_029 | PunanTubu | 6 | 6.6 | 1.5 | 55 | F | 2 | 50.54 | 2.67 | 20.18 | 9.72 | 3.41 | 7.68 |
| MAL_RPNII_030 | PunanTubu | 7 | 7.4 | 3.6 | 69 | M | 1 | 50.22 | 2.06 | 17.26 | 9.34 | 5.65 | 7.28 |
| MAL_RPNII_033 | PunanTubu | 4 | 6.1 | 1 | 44 | M | 2 | 51.84 | 2.27 | 20.52 | 7.79 | 4.71 | 6.78 |
| MAL_RPNII_035 | PunanTubu | 3 | 6.8 | 2.1 | 35 | M | 2 | 49.34 | 2.68 | 21.09 | 9.61 | 5.59 | 6.71 |
| MAL_RPNII_036 | PunanTubu | 2 | 7 | 1.4 | 60 | F | 2 | 51.21 | 2.57 | 21.67 | 8.26 | 4.21 | 7.13 |
| MAL_RPNII_038 | PunanTubu | 5 | 5.5 | 1.1 | 32 | M | 1 | 51.87 | 2.30 | 18.92 | 9.31 | 4.12 | 6.97 |
| MAL_RPNII_039 | PunanTubu | 6 | 7.4 | 1.3 | 49 | F | 2 | 46.11 | 2.87 | 25.24 | 10.59 | 4.83 | 6.25 |
| MAL_RPNII_040 | PunanTubu | 3 | 5.7 | 0.8 | 52 | M | 1 | 48.12 | 1.90 | 19.61 | 11.05 | 5.11 | 6.61 |
| MAL_RPNII_042 | PunanTubu | 1 | 5.5 | 0.6 | 29 | F | 2 | 55.45 | 2.77 | 19.55 | 8.30 | 3.84 | 6.73 |
| MAL_SPIII_001 | Lundayeh | 2 | 6.2 | 1.5 | 69 | M | 2 | 50.37 | 2.06 | 19.47 | 8.95 | 4.85 | 6.84 |
| MAL_SPIII_002 | Lundayeh | 1 | 7 | 1.5 | 55 | F | 1 | 52.36 | 2.78 | 22.61 | 9.37 | 3.58 | 6.07 |
| MAL_SPIII_003 | Lundayeh | 3 | 6.6 | 1.1 | 67 | M | 2 | 52.45 | 2.35 | 20.52 | 11.22 | 4.33 | 6.52 |
| MAL_SPIII_004 | Lundayeh | 2 | 6 | 1.6 | 69 | F | 1 | 48.65 | 2.31 | 23.96 | 11.37 | 4.77 | 6.51 |
| MAL_SPIII_005 | Lundayeh | 6 | 7.2 | 1.5 | 72 | M | 2 | 44.67 | 2.98 | 21.36 | 8.77 | 5.96 | 7.04 |
| MAL_SPIII_006 | Lundayeh | 1 | 6.9 | 1.6 | 47 | M | 2 | 47.09 | 2.55 | 21.56 | 9.55 | 5.26 | 6.87 |
| MAL_SPIII_007 | Lundayeh | 5 | 5.4 | 1.2 | 40 | M | 1 | 47.64 | 2.64 | 21.31 | 9.76 | 4.52 | 6.63 |
| MAL_SPIII_008 | Lundayeh | 3 | 7.5 | 1.5 | 47 | F | 1 | 46.77 | 2.76 | 24.97 | 10.53 | 4.23 | 6.33 |
| MAL_SPIII_009 | Lundayeh | 6 | 6.9 | 1.3 | 67 | M | 2 | 50.71 | 2.05 | 21.69 | 7.83 | 4.26 | 7.09 |
| MAL_SPIII_010 | Lundayeh | 4 | 6.1 | 1.6 | 37 | M | 1 | 49.20 | 1.98 | 18.46 | 8.29 | 4.63 | 6.62 |
| MAL_SPIII_011 | Lundayeh | 3 | 6.1 | 1.4 | 44 | M | 2 | 49.61 | 2.42 | 20.28 | 10.39 | 4.92 | 6.39 |
| MAL_SPIII_012 | Lundayeh | 5 | 7 | 1.3 | 55 | M | 2 | 50.21 | 2.25 | 21.40 | 9.37 | 4.48 | 7.10 |
| MAL_SPIII_013 | Lundayeh | 3 | 5.9 | 1.3 | 57 | M | 2 | 49.78 | 2.79 | 20.65 | 8.54 | 4.80 | 6.66 |
| MAL_SPIII_014 | Lundayeh | 6 | 6.6 | 1.7 | 31 | F | 1 | 50 | 2.87 | 22.69 | 8.94 | 5.05 | 6.37 |
| MAL_SPIII_015 | Lundayeh | 6 | 6.6 | 1.2 | 27 | F | 2 | 46.88 | 2.59 | 22.11 | 10.86 | 4.32 | 6.46 |
| MAL_SPIII_017 | Lundayeh | 5 | 5.8 | 1 | 36 | M | 2 | 44.39 | 2.77 | 21.93 | 10.05 | 5.54 | 6.63 |
| MAL_SPIII_023 | Lundayeh | 7 | 7.7 | 2.8 | 28 | F | 2 | 47.85 | 2.97 | 21.83 | 9.97 | 4.84 | 7.04 |
| MAL_SPIII_019 | Lundayeh | 2 | 6.5 | 1.5 | 61 | F | 1 | 47.29 | 2.66 | 21.88 | 9.70 | 3.79 | 6.89 |
| MAL_SPIII_020 | Lundayeh | 4 | 7.3 | 2.6 | 70 | M | 1 | 50.97 | 2.42 | 17.81 | 8.49 | 5.32 | 7.39 |
| MAL_SPIII_022 | Lundayeh | 5 | 6.4 | 1.6 | 62 | M | 1 | 51.77 | 2.40 | 17.65 | 7.63 | 4.56 | 6.91 |
| MAL_SPIII_036 | Lundayeh | 7 | 7.3 | 1.6 | 55 | M | 1 | 48.04 | 2.59 | 21.87 | 10.53 | 3.75 | 6.80 |
| MAL_SPIII_024 | Lundayeh | 4 | 6.4 | 1.9 | 72 | M | 1 | 49.42 | 1.79 | 18.20 | 8.39 | 6.16 | 6.86 |
| MAL_SPIII_025 | Lundayeh | 6 | 6.40 | 1.2 | 50 | F | 2 | 50.20 | 2.83 | 20.74 | 9.93 | 3.91 | 6.77 |
| MAL_SPIII_027 | Lundayeh | 1 | 6.2 | 0.9 | 59 | M | 2 | 54.58 | 2.34 | 17.83 | 8.02 | 4.57 | 7.15 |
| MAL_SPIII_028 | Lundayeh | 6 | 7.1 | 1.7 | 68 | F | 2 | 48.38 | 2.82 | 22.70 | 9.21 | 5.15 | 6.53 |
| MAL_SPIII_030 | Lundayeh | 1 | 6.9 | 1.3 | 71 | F | 2 | 45.35 | 2.33 | 21.57 | 9.82 | 5.75 | 6.70 |

| | | | | | | | | | | | | | |
|---|---|---|---|---|---|---|---|---|---|---|---|---|---|
| MAL_SPIII_031 | Lundayeh | 4 | 6.7 | 1.8 | 46 | F | 1 | 49.36 | 2.46 | 20.60 | 9.27 | 5.10 | 7.05 |
| MAL_SPIII_032 | Lundayeh | 1 | 7.1 | 1.7 | 54 | F | 1 | 54.27 | 2.51 | 21.64 | 7.63 | 3.08 | 6.93 |
| MAL_SPIII_033 | Lundayeh | 1 | 4.9 | 0.6 | 46 | F | 2 | 50.50 | 2.69 | 21.68 | 7.98 | 4.95 | 6.51 |
| MAL_SPIII_034 | Lundayeh | 5 | 7.2 | 1.4 | 36 | F | 2 | 51.85 | 2.42 | 20.64 | 9.43 | 3.92 | 6.78 |
| MAL_SPIII_018 | Lundayeh | 7 | 7.1 | 1.4 | 65 | F | 2 | 48.54 | 2.11 | 17.46 | 9.06 | 5.41 | 6.91 |
| MAL_SPIII_037 | Lundayeh | 2 | 6.3 | 1.6 | 40 | M | 1 | 50.37 | 2.24 | 21.05 | 9.89 | 3.67 | 6.88 |
| MAL_SPIII_038 | Lundayeh | 2 | 7.5 | 1.7 | 27 | F | 2 | 47.60 | 2.86 | 21.49 | 11.85 | 5.05 | 6.30 |
| MAL_SPIII_039 | Lundayeh | 3 | 6.7 | 1.7 | 36 | M | 2 | 47.76 | 2.59 | 22.89 | 11.25 | 5.38 | 6.51 |

**Table S2. RNA Seq data covariates.**

| variable | statistic | p | p.adj | effsize | magnitude | method | groups |
|---|---|---|---|---|---|---|---|
| Bio_VisceralFat | 30.82104 | 2.03E-07 | 1.83E-06 | 0.27713 | large | Kruskal-Wallis | PBvPTvLDY |
| Bio_BMRI | 27.00135 | 1.37E-06 | 1.23E-05 | 0.24040 | large | Kruskal-Wallis | PBvPTvLDY |
| Bio_BMI | 24.30796 | 5.27E-06 | 4.74E-05 | 0.21450 | large | Kruskal-Wallis | PBvPTvLDY |
| Bio_BodyFat | 15.33960 | 4.67E-04 | 0.004203 | 0.12827 | moderate | Kruskal-Wallis | PBvPTvLDY |
| Blood_Cholesterol | 12.25739 | 0.00218 | 0.01962 | 0.09863 | moderate | Kruskal-Wallis | PBvPTvLDY |
| Blood_Glucose | 9.75948 | 0.0076 | 0.0684 | 0.07461 | moderate | Kruskal-Wallis | PBvPTvLDY |
| Blood_UricAcid | 9.40245 | 0.00908 | 0.08172 | 0.07118 | moderate | Kruskal-Wallis | PBvPTvLDY |
| Bio_SubcutFat_Overall | 9.33736 | 0.00938 | 0.08442 | 0.07055 | moderate | Kruskal-Wallis | PBvPTvLDY |
| Blood_LDL | 7.66402 | 0.0217 | 0.1953 | 0.05446 | small | Kruskal-Wallis | PBvPTvLDY |
| Bio_VisceralFat | 550 | 2.72E-05 | 2.45E-04 | 0.40598 | moderate | Wilcoxon | HGvAGR |
| Bio_BMI | 572.5 | 5.41E-05 | 4.87E-04 | 0.39063 | moderate | Wilcoxon | HGvAGR |
| Bio_BodyFat | 611 | 1.64E-04 | 0.001476 | 0.36475 | moderate | Wilcoxon | HGvAGR |
| Blood_Cholesterol | 658 | 5.74E-04 | 0.005166 | 0.33326 | moderate | Wilcoxon | HGvAGR |
| Blood_Triglyceride | 676.5 | 9.15E-04 | 0.008235 | 0.32086 | moderate | Wilcoxon | HGvAGR |
| Bio_SubcutFat_Overall | 717.5 | 0.00244 | 0.02196 | 0.29334 | small | Wilcoxon | HGvAGR |
| Blood_Glucose | 731.5 | 0.00334 | 0.03006 | 0.28403 | small | Wilcoxon | HGvAGR |
| Blood_LDL | 756.5 | 0.00576 | 0.05184 | 0.26724 | small | Wilcoxon | HGvAGR |
| Bio_BMRI | 793 | 0.0122 | 0.1098 | 0.24272 | small | Wilcoxon | HGvAGR |

**Table S3. Summary of biological and blood variables across different group comparisons.**

| Task | Model | Accuracy | Kappa | Sensitivity / Balanced Accuracy |
|---|---|---|---|---|
| **Ternary** | LDA | 47.5% | 0.22 | PB: 66.7% / 76.2%<br>PT: 57.1% / 74.7%<br>LDY: 85.7% / 81.3% |
| | QDA | 53.1% | 0.30 | PB: 76.7% / 76.0%<br>PT: 33.3% / 53.4%<br>LDY: 55.3% / 68.2% |
| | RF | 56.1% | 0.34 | PB: 60.0% / 69.6%<br>PT: 53.9% / 62.2%<br>LDY: 55.3% / 69.7% |
| **Binary** | LDA | 70.1% | 0.23 | HG: 40.0% / 60.9% |
| | QDA | 68.2% | 0.33 | HG: 73.3% / 69.8% |
| | RF | 76.6% | 0.40 | HG: 53.3% / 69.5% |

**Table S4. Classification performance for ternary (PB vs. PT vs. LDY) and binary (HG vs. AGR) tasks using 5-fold cross-validation.**

| ID | Description | setSize | enrichScr | NES | pvalue | p.adjust | qvalue | compar |
|---|---|---|---|---|---|---|---|---|
| GO:0002181 | cytoplasmic translation | 149 | -0.61 | -2.21 | 0.00 | 0.00 | 0.00 | PBvLDY |
| GO:0090329 | regulation of DNA-templated DNA replication | 41 | 0.75 | 2.20 | 0.00 | 0.00 | 0.00 | PBvLDY |
| GO:0086004 | regulation of cardiac muscle cell contraction | 16 | 0.85 | 2.05 | 0.00 | 0.01 | 0.01 | PBvLDY |
| GO:0006260 | DNA replication | 245 | 0.44 | 1.69 | 0.00 | 0.01 | 0.01 | PBvLDY |
| GO:1903115 | regulation of actin filament-based movement | 18 | 0.82 | 2.03 | 0.00 | 0.02 | 0.02 | PBvLDY |
| GO:0002250 | adaptive immune response | 361 | 0.40 | 1.58 | 0.00 | 0.02 | 0.02 | PBvLDY |
| GO:1902107 | positive regulation of leukocyte differentiation | 147 | 0.48 | 1.71 | 0.00 | 0.03 | 0.03 | PBvLDY |
| GO:1903708 | positive regulation of hemopoiesis | 147 | 0.48 | 1.71 | 0.00 | 0.03 | 0.03 | PBvLDY |
| GO:0006275 | regulation of DNA replication | 108 | 0.52 | 1.79 | 0.00 | 0.03 | 0.03 | PBvLDY |
| GO:0050727 | regulation of inflammatory response | 252 | 0.41 | 1.56 | 0.00 | 0.04 | 0.03 | PBvLDY |
| GO:0002922 | positive regulation of humoral immune response | 10 | 0.88 | 1.88 | 0.00 | 0.04 | 0.04 | PBvLDY |
| GO:0006261 | DNA-templated DNA replication | 144 | 0.47 | 1.69 | 0.00 | 0.04 | 0.04 | PBvLDY |
| GO:0051276 | chromosome organization | 496 | 0.35 | 1.45 | 0.00 | 0.05 | 0.04 | PBvLDY |
| GO:0002675 | positive regulation of acute inflammatory response | 18 | 0.79 | 1.96 | 0.00 | 0.05 | 0.04 | PBvLDY |
| GO:0009615 | response to virus | 310 | 0.38 | 1.48 | 0.00 | 0.05 | 0.04 | PBvLDY |
| GO:0002521 | leukocyte differentiation | 478 | 0.35 | 1.45 | 0.00 | 0.05 | 0.04 | PBvLDY |
| GO:0002181 | cytoplasmic translation | 149 | -0.53 | -1.93 | 0.00 | 0.00 | 0.00 | PBvPT |
| GO:0051276 | chromosome organization | 496 | 0.37 | 1.53 | 0.00 | 0.02 | 0.02 | PBvPT |

**Table S5. GSEA GO:BP Differential Expression.**

| ID | Description | GeneRat | BgRat | FoldEnric | zScore | pval | p.adj | qval | compar |
|---|---|---|---|---|---|---|---|---|---|
| GO:0003013 | circulatory system process | 30/355 | 315/10826 | 2.90 | 6.32 | 0.00 | 0.00 | 0.00 | PBvLDY |
| GO:0007186 | G protein-coupled receptor signaling pathway | 32/355 | 363/10826 | 2.69 | 6.02 | 0.00 | 0.00 | 0.00 | PBvLDY |
| GO:0010885 | regulation of cholesterol storage | 6/355 | 14/10826 | 13.07 | 8.32 | 0.00 | 0.00 | 0.00 | PBvLDY |
| GO:0008015 | blood circulation | 24/355 | 259/10826 | 2.83 | 5.48 | 0.00 | 0.00 | 0.00 | PBvLDY |
| GO:0010742 | macrophage derived foam cell differentiation | 7/355 | 23/10826 | 9.28 | 7.32 | 0.00 | 0.00 | 0.00 | PBvLDY |
| GO:0090077 | foam cell differentiation | 7/355 | 24/10826 | 8.89 | 7.13 | 0.00 | 0.00 | 0.00 | PBvLDY |
| GO:0009617 | response to bacterium | 31/355 | 409/10826 | 2.31 | 4.98 | 0.00 | 0.01 | 0.01 | PBvLDY |
| GO:0042742 | defense response to bacterium | 17/355 | 157/10826 | 3.30 | 5.35 | 0.00 | 0.01 | 0.01 | PBvLDY |
| GO:0010743 | regulation of macrophage derived foam cell diff. | 6/355 | 18/10826 | 10.17 | 7.17 | 0.00 | 0.01 | 0.01 | PBvLDY |
| GO:0010878 | cholesterol storage | 6/355 | 19/10826 | 9.63 | 6.93 | 0.00 | 0.01 | 0.01 | PBvLDY |
| GO:0097006 | regulation of plasma lipoprotein particle levels | 8/355 | 42/10826 | 5.81 | 5.75 | 0.00 | 0.02 | 0.02 | PBvLDY |
| GO:0009636 | response to toxic substance | 15/355 | 151/10826 | 3.03 | 4.62 | 0.00 | 0.04 | 0.03 | PBvLDY |
| GO:0032613 | interleukin-10 production | 8/355 | 48/10826 | 5.08 | 5.22 | 0.00 | 0.04 | 0.03 | PBvLDY |
| GO:0032653 | regulation of interleukin-10 production | 8/355 | 48/10826 | 5.08 | 5.22 | 0.00 | 0.04 | 0.03 | PBvLDY |
| GO:0002385 | mucosal immune response | 5/355 | 17/10826 | 8.97 | 6.05 | 0.00 | 0.04 | 0.03 | PBvLDY |
| GO:0070663 | regulation of leukocyte proliferation | 17/355 | 191/10826 | 2.71 | 4.40 | 0.00 | 0.04 | 0.03 | PBvLDY |
| GO:0002251 | organ or tissue specific immune response | 5/355 | 18/10826 | 8.47 | 5.84 | 0.00 | 0.04 | 0.04 | PBvLDY |

**Table S6. ORA GO:BP Differential Expression.**

| TERM | N | DE | P.DE | FDR | compar |
|---|---|---|---|---|---|
| cell morphogenesis | 966.00 | 384.00 | 0.00 | 0.03 | PBvLDY |
| system process | 2208.00 | 664.00 | 0.00 | 0.05 | PBvLDY |
| catecholamine metabolic process | 54.00 | 29.00 | 0.00 | 0.05 | PBvLDY |
| cell adhesion | 1511.00 | 522.00 | 0.00 | 0.03 | PBvLDY |
| anatomical structure morphogenesis | 2730.00 | 924.00 | 0.00 | 0.03 | PBvLDY |
| catechol-containing compound metabolic process | 54.00 | 29.00 | 0.00 | 0.05 | PBvLDY |
| dopamine metabolic process | 39.00 | 25.00 | 0.00 | 0.01 | PBvLDY |
| sensory perception of mechanical stimulus | 183.00 | 83.00 | 0.00 | 0.05 | PBvLDY |
| response to stimulus | 8596.00 | 4575.00 | 0.00 | 0.00 | PBvPT |
| multicellular organismal process | 7641.00 | 4228.00 | 0.00 | 0.00 | PBvPT |
| cellular response to stimulus | 7270.00 | 3860.00 | 0.00 | 0.00 | PBvPT |
| anatomical structure development | 5912.00 | 3351.00 | 0.00 | 0.00 | PBvPT |
| signaling | 6291.00 | 3467.00 | 0.00 | 0.00 | PBvPT |
| signal transduction | 5805.00 | 3169.00 | 0.00 | 0.00 | PBvPT |
| developmental process | 6443.00 | 3585.00 | 0.00 | 0.00 | PBvPT |
| cellular developmental process | 4349.00 | 2485.00 | 0.00 | 0.00 | PBvPT |
| cell differentiation | 4347.00 | 2483.00 | 0.00 | 0.00 | PBvPT |
| system development | 4014.00 | 2337.00 | 0.00 | 0.00 | PBvPT |
| response to chemical | 3916.00 | 2083.00 | 0.00 | 0.00 | PBvPT |
| regulation of cell communication | 3497.00 | 1969.00 | 0.00 | 0.00 | PBvPT |
| regulation of signaling | 3492.00 | 1960.00 | 0.00 | 0.00 | PBvPT |
| regulation of multicellular organismal process | 3085.00 | 1704.00 | 0.00 | 0.00 | PBvPT |
| animal organ development | 2994.00 | 1775.00 | 0.00 | 0.00 | PBvPT |
| cell development | 2807.00 | 1637.00 | 0.00 | 0.00 | PBvPT |
| regulation of biological quality | 2891.00 | 1641.00 | 0.00 | 0.00 | PBvPT |
| cell surface receptor signaling pathway | 2768.00 | 1594.00 | 0.00 | 0.00 | PBvPT |
| nervous system development | 2503.00 | 1466.00 | 0.00 | 0.00 | PBvPT |
| immune system process | 2450.00 | 1297.00 | 0.00 | 0.04 | PBvPT |
| intracellular signal transduction | 2682.00 | 1503.00 | 0.00 | 0.02 | PBvPT |
| cellular response to chemical stimulus | 2683.00 | 1485.00 | 0.00 | 0.00 | PBvPT |
| multicellular organism development | 4698.00 | 2683.00 | 0.00 | 0.00 | PBvPT |
| transport | 4302.00 | 2367.00 | 0.00 | 0.00 | PBvPT |
| establishment of localization | 4604.00 | 2495.00 | 0.00 | 0.02 | PBvPT |
| cell communication | 6383.00 | 3522.00 | 0.00 | 0.00 | PBvPT |
| response to external stimulus | 2663.00 | 1459.00 | 0.00 | 0.00 | PBvPT |
| positive regulation of response to stimulus | 2299.00 | 1280.00 | 0.00 | 0.00 | PBvPT |
| positive regulation of cell communication | 1790.00 | 1037.00 | 0.00 | 0.00 | PBvPT |
| positive regulation of signaling | 1791.00 | 1035.00 | 0.00 | 0.00 | PBvPT |
| cell motility | 1794.00 | 1062.00 | 0.00 | 0.00 | PBvPT |
| neurogenesis | 1728.00 | 1050.00 | 0.00 | 0.00 | PBvPT |
| cell-cell signaling | 1702.00 | 1049.00 | 0.00 | 0.00 | PBvPT |
| immune response | 1670.00 | 900.00 | 0.00 | 0.00 | PBvPT |
| positive regulation of multicellular organismal process | 1668.00 | 957.00 | 0.00 | 0.00 | PBvPT |
| generation of neurons | 1495.00 | 924.00 | 0.00 | 0.00 | PBvPT |
| response to biotic stimulus | 1507.00 | 785.00 | 0.00 | 0.04 | PBvPT |
| transmembrane transport | 1530.00 | 896.00 | 0.00 | 0.00 | PBvPT |
| cell projection organization | 1597.00 | 968.00 | 0.00 | 0.00 | PBvPT |
| plasma membrane bounded cell projection organization | 1555.00 | 940.00 | 0.00 | 0.00 | PBvPT |
| cell migration | 1584.00 | 933.00 | 0.00 | 0.00 | PBvPT |
| positive regulation of signal transduction | 1576.00 | 903.00 | 0.00 | 0.01 | PBvPT |
| monoatomic ion transport | 1239.00 | 751.00 | 0.00 | 0.00 | PBvPT |

**Table S7. Enrichment analyses using `gometh()` GO:BP Differential Methylation. PBvPT only showing top-50 enriched terms.**

| Model | Free parameters | $k$ |
|---|---|---|
| BM | $\sigma^2,\ \mu_0,\ V_{\text{within},i}$ | 5 |
| $\text{OU}_{\text{shared}}$ | $+\ \alpha,\ \theta$ | 6 |
| $\text{OU}_{\theta}$ | $+\ \alpha,\ \theta_{PB},\ \theta_{PT},\ \theta_{LDY}$ | 8 |
| $\text{OU}_{\alpha}$ | $+\ \alpha_{PB},\ \alpha_{PT},\ \alpha_{LDY},\ \theta$ | 8 |
| $\text{OU}_{\alpha\theta}$ | $+\ \alpha_{PB},\ \alpha_{PT},\ \alpha_{LDY},\ \theta_{PB},\ \theta_{PT},\ \theta_{LDY}$ | 10 |

**Table S8. OU models fitted per gene.**
